# Model-driven engineering of *Yarrowia lipolytica* for improved microbial oil production

**DOI:** 10.1101/2024.07.31.606002

**Authors:** Zeynep Efsun Duman-Özdamar, Mattijs K. Julsing, Vitor A.P. Martins dos Santos, Jeroen Hugenholtz, Maria Suarez-Diez

## Abstract

Extensive usage of plant-based oils, especially palm oil, has led to environmental and social issues, such as deforestation and loss of biodiversity, thus sustainable alternatives are required. Microbial oils, especially from *Yarrowia lipolytica*, offer a promising solution due to their similar composition to palm oil, low carbon footprint, and ability to utilize low-cost substrates. In this study, we employed the Design-Build-Test-Learn (DBTL) approach to enhance lipid production in *Y. lipolytica*. We systematically evaluated predictions from the genome-scale metabolic model to identify and overcome bottlenecks in lipid biosynthesis. We tested the effect of predicted medium supplements and genetic intervention targets, including the overexpression of ATP-citrate lyase (*ACL*), acetyl-CoA carboxylase (*ACC*), threonine synthase (*TS*), diacylglycerol acyltransferase(*DGA1*), the deletion of citrate exporter gene (*CEX1*) and disruption of β-oxidation pathway (*MFE1*). Combining *TS* and *DGA1* overexpression in the *Δmfe_Δcex* background achieved a remarkable 200% increase in lipid content (56 % w/w) and a 230% increase in lipid yield on glycerol. These findings underscore the potential of *Y. lipolytica* as an efficient microbial cell factory for fatty acid production. Our study advances the understanding of lipid metabolism in *Y. lipolytica* and demonstrates a viable approach for developing sustainable and economically feasible alternatives to palm oil.

**Graphical Abstract:** 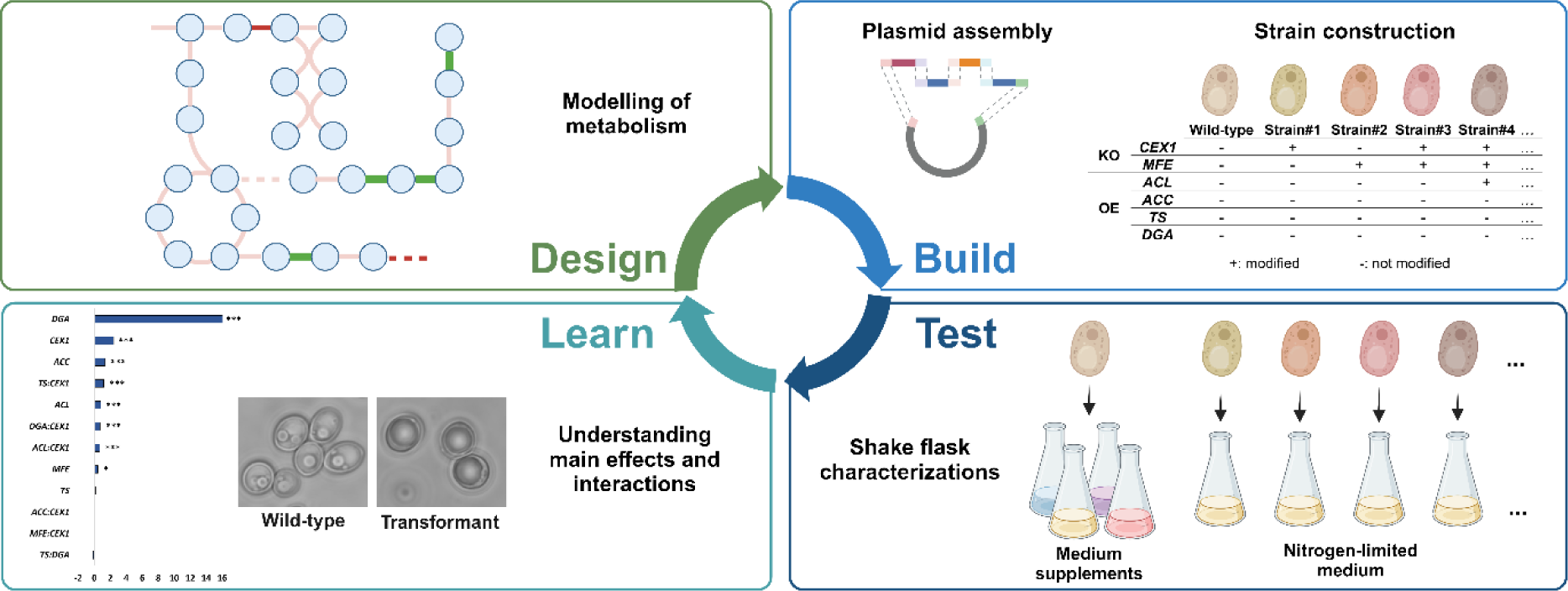

We followed the Design-Build-Test-Learn approach to identify and overcome bottlenecks in lipid biosynthesis in *Y. lipolytica*. DBTL intertwined the predictions from the metabolic model with addressed bottlenecks, investigated the effect of genetic interventions and medium supplements on lipid content, and ultimately defined an efficient strain design strategy.

## Introduction

Plant-based oils are extensively used in food, feed, chemical, personal care, and cosmetic products to enhance texture, flavor, and shelf-life (Holley and Patel, 2005; P. Desbois, 2012). Palm oil, in particular, is favored as an inexpensive source of these functional components (Rustan and Drevon, 2005). However, the rising demand for palm oil has led to the destruction of native tropical forests in many countries across Asia, South America, and Africa, has severe consequences for local communities and contributes to climate change (Vijay et al., 2016; Abubakar et al., 2021; Murphy et al., 2021). Therefore, there is an urgent and critical need to develop sustainable alternatives to palm-based fatty acids and oils.

Microbial oils present a valuable alternative to traditional plant-based oils due to their sustainability and versatility. They can be produced using renewable resources and waste substrates, minimizing ecological impact and promoting circular economy practices. Among the oil-producing microorganisms, oleaginous yeasts contain more than 20% of their total biomass in lipids (Salvador López et al., 2022). Oleaginous yeasts are often considered superior for commercial applications due to their fast growth, high lipid content, and high volumetric productivity (Sitepu et al., 2014). The most extensively studied oleaginous yeast have been *Cutaneotrichosporon oleaginosus*, *Rhodotorula toruloides*, and *Yarrowia lipolytica* (Abeln and Chuck, 2021).

*Yarrowia lipolytica* commonly accumulates lipids up to 20-30 % of its biomass under nitrogen-limiting conditions (Beopoulos et al., 2009). Moreover, *Y. lipolytica* is non-pathogenic and regarded as food-grade yeast, thus its oil can be used for food-related applications (Zinjarde, 2014; Amalia *et al*., 2020). Under nitrogen-limiting conditions, *Y. lipolytica* produces fatty acids comparable to that of palm, composed of 15 % palmitic acid (C16:0), 13% stearic acid (C18:0), 51% oleic acid (C18:1), and 21% linoleic acid (C18:2) (Carsanba *et al*., 2018). It is able to grow and produce lipids on a wide range of substrates including agro-industrial residues, and crude glycerol (Papanikolaou and Aggelis, 2003; Rywińska *et al*., 2013; Sara *et al*., 2016; Caporusso *et al*., 2021). Due to these advantages, this oleaginous yeast is flagged as an attractive microbial-cell factory to sustain a bio-based circular economy for industrial implementation.

Lipid synthesis in *Y. lipolytica* occurs via *de novo* synthesis by metabolizing hydrophilic substrates (A. Fabiszewska *et al*., 2019). When nitrogen is limited in the medium, the excess carbon is directed to fatty acid synthesis via channeling mitochondrial citrate to the cytoplasm. ATP citrate lyase (*ACL*) converts citrate into acetyl-CoA and oxaloacetate then acetyl-CoA carboxylase (*ACC*) converts acetyl-CoA to malonyl-CoA which is the primary precursor for fatty acid elongation (Fabiszewska *et al*., 2019; Poontawee *et al*., 2023). Followed by fatty acid elongation, synthesized acyl-CoA chains are finally incorporated into triacylglycerol (TAG) by diacylglycerol acyltransferase (*DGA1*).

Several research groups worked on strategies for increasing lipid accumulation in *Y. lipolytica* and developed metabolic engineering tools and strategies mainly focused on maximizing the flux toward lipid biosynthesis (Larroude *et al*., 2019; Wang *et al*., 2020). Tai and Stephanopoulos, 2013 explored the push and pull strategy for lipid accumulation by overexpression of *ACC1* and *DGA1.* Overexpression of *ACL* from *Mus musculus* in *Y. lipolytica* enhanced the citrate conversion to acetyl-CoA resulting in higher lipid contents (Zhang *et al*., 2014). Furthermore minimizing the flux toward one of the competing metabolic pathways was achieved by the deletion of genes related to the β-oxidation pathway, which degrades the intracellular fatty acids, (Dulermo and Nicaud, 2011; Blazeck *et al*., 2013). In addition to these strategies, (Kim et al., 2019) analyzed the genome-scale metabolic model (GEM) of *Y. lipolytica* and successfully predicted some of the established genetic engineering strategies for higher lipid accumulation. Although the main limitations leading to lower lipid accumulation levels in *Y. lipolytica* were highlighted over the last two decades, these bottlenecks have not been addressed systematically.

In this study, we followed the Design-Build-Test-Learn (DBTL) approach, a streamlined method for iterating the steps of strain development. This approach integrates systems biology and metabolic engineering to develop *Y. lipolytica* into a sustainable and more productive fatty acid production platform. We intertwined the predictions from the GEM of *Y. lipolytica* with previously addressed bottlenecks for improved lipid accumulation and ultimately defined an efficient strain design strategy. The identified genetic intervention strategy was experimentally validated with iterations in the built and test step.

## Experimental Procedures

### 1. Comparative flux sampling analysis using the GEM

We used Comparative Flux Sampling Analysis (CFSA) to identify suitable strategies by using the iYali4, v4.1.2 model (Kerkhoven et al., 2016; van Rosmalen et al., 2024). In brief, the model was used to simulate scenarios of maximum growth, maximum lipid production, and slow growth. In each scenario flux sampling was used to characterize the metabolic space and reactions with the highest changes between scenarios were selected as targets for further inspection as described by (van Rosmalen et al., 2024). The N-limiting medium was adjusted by using glycerol exchange (y001808) and urea exchange (y002091) reactions. CFSA was performed (number of samples= 30000, optimality = 0.90, flux fraction = 1.25, KS1 = KS2 ≥ 0.75, mean absolute change ≥ 0.01, standard deviation in production ≤ 50) using the lipid exchange reaction (xlipid_export) as a target for the production scenario and xBIOMASS for growth scenario. CFSA is available at GitLab.

### 2. Strains, media, and growth conditions in shake-flask

*Yarrowia lipolytica* strains used in this study were derived from wild-type *Y. lipolytica* CBS8108 strain obtained from Westerdijk Fungalbio Diversity Institute (Utrecht, The Netherlands) and maintained on Yeast extract Peptone Dextrose (YPD) agar plates containing 10 g/L yeast extract, 20 g/L peptone, 20 g/L glucose, 20 g/L agar. The maintained cultures were stored at 4 °C for up to a week. *Escherichia coli* Zymo 10B (Zymo Research, Orange, CA) was used for all cloning purposes throughout this study and maintained on Luria-Bertani (LB) agar (10 g/L tryptone, 10 g/L NaCl, 5 g/L yeast extract, 15 g/L agar) with ampicillin (100 μg/ml) at 37°C.

The inoculum was prepared by transferring a single colony of *Y. lipolytica* into 10 mL YPD broth (10 g/L yeast extract, 20 g/L peptone, 20 g/L glucose, 20 g/L) in 50 mL tubes and incubated at 30 °C, 250 rpm for 18 h in a shaking incubator. Wild-type and other built *Y. lipolytica* transformants were cultivated into minimal media consisting of glycerol as carbon source and urea as nitrogen source with set ratios of C/N (g/g) 140 (Duman-Özdamar et al., 2022). Methionine (2 mM), threonine (2 mM), leucine (2 mM), and glutamate (2 mM) were added into C/N 140 cultivation medium, and only wild-type was tested in these experiments. Cultures were incubated at 30 °C, 250 rpm for 120 h in a shaking incubator. Cells were harvested at the end of incubation and centrifuged at 1780 g, 4 °C for 20 min. All experiments were performed in triplicates.

### 3. Plasmid construction and preparation for transformation

Restriction enzymes and Q5 High-Fidelity DNA polymerase used in cloning were obtained from New England Biolabs (Ipswich, MA). Genomic DNA (gDNA) of *Y. lipolytica* was prepared using YeaStar Genomic DNA Kit (Zymo Research, Irvine, CA). PCR products and DNA fragments were purified with GeneJET Gel Extraction Kit (Thermo Scientific, Waltham, MA). The primers and plasmids used are described in Table S1 and Table S2 respectively. Assembly of the plasmids was performed by using NEBridge Golden Gate Assembly Kit (BsaI-HF v2) (New England Biolabs, Ipswich, MA). All constructed plasmids were verified by whole plasmid sequencing (Eurofins, Germany).

ATP-citrate lyase (*ACL1)*, acetyl-CoA carboxylase containing introns (*ACC)*, and threonine synthase (*TS)* genes from *C. oleaginosus* were amplified from pUC57-ACL, pUC57-ACC, pUC57-TS plasmid (Duman-Özdamar et al., 2024) by using primers ACL_BsaI_D_Fw and ACL_BsaI_E_Rv, ACC_BsaI_D_Fw and ACC_BsaI_E_Rv, TS_BsaI_D_Fw and TS_BsaI_E_Rv respectively (Table S1). Diacylglycerol acyltransferase 1 gene (*DGA1,* Accession Number: XM_504700) and homologous upstream and downstream parts of the citrate exporter gene of *Y. lipolytica* (*CEX1,* Accession Number: XM_503062.1) were amplified from the gDNA by using primers DGA_D_BsaI_Fw and DGA_E_BsaI_Rv, Cex1Up_BsaI_A_Fw and Cex1Up_BsaI_B_Rv, Cex1Down_BsaI_C_Fw and Cex1Down_BsaI_M_Rv, Cex1Down_BsaI_L_Fw and Cex1Down_BsaI_M_Rv. Amplified parts were cloned into the pCR-Blunt vector using Zero Blunt™ PCR Cloning Kit by following the instructions from the supplier (Invitrogen, Waltham, MA). These amplified parts were assembled with the parts from the *Yarrowia lipolytica* Golden Gate tool kit (Addgene kit #1000000167) by facilitating BsaI restriction sites; promoter (pCR4Blunt-TOPO- P1 TEF-8UAS), terminator (pCR4Blunt-TOPO-TLip2 (E-L)), two markers for antibiotic resistance (hygromycin, pCR4Blunt-TOPO-M-hph, and nourseothricin, pCR4Blunt-TOPO-M-Nat), MFE homologous sites with NotI restriction sites (pCR4Blunt-TOPO-MFE-NotI_Up, pCR4Blunt- TOPO-MFE-NotI_Down) and backbone plasmid (pSB1A3) (Larroude et al., 2019).

Assembled plasmids were transformed into *E. coli* Zymo 10B cells (Cat #T3020; Zymo Research, Irvine, CA, The US) by following the supplier’s instructions, and the transformed strains were stored at -80 °C. *E. coli* cells were grown overnight in 10 ml LB broth with 100µg/ml ampicillin or 50 µg/ml kanamycin in 50 mL falcon tubes shaking at 250 rpm at 37 °C. Plasmid DNA was isolated using the GeneJET Plasmid Miniprep Kit (Cat #K0503; Thermo Fisher Scientific, MA, The US). Isolated plasmids were prepared for transformation by releasing the assembled cassette cut with NotI restriction enzyme according to the supplier’s instructions (NEB, Ipswich, MA, The US).

### 4. Transformation and selection for transformants

Electrocompetent *Y. lipolytica* cells were prepared by following the protocol established by (Duman-Özdamar et al., 2024). Approximately 1 µg linearized vector was mixed with 50 µl electrocompetent *Y. lipolytica* cells and incubated on ice for 5 minutes. Electroporation was performed using a pulse of 0.8 kVolt, 1000 Ohm, 25 µFarad (Bio-Rad, CA, The US) in 2 mm electroporation cuvettes. 1 mL of YPD broth was added immediately after pulsing. The cells were transferred to a 2 mL Eppendorf tube, and incubated for 2.5 hours at 30 °C, gently mixing the cells every 30 minutes by inversion. 100 µL cells were spread onto YPD agar plate containing 300 µg/mL nourseothricin and/or 200 µg/mL hygromycin for primary selection. The negative control was plated on YPD agar. Incubation of the plates was done at 30 °C for 48 hours. Grown colonies were randomly selected and streaked into a YPD agar plate including 400 µg/mL nourseothricin and/or 300 µg/mL hygromycin for secondary selection. Transformants were confirmed via colony PCR. The primers in Table S1 and DreamTaq Green PCR Master Mix (2X) were used by following the instructions from the supplier (Thermo Fisher Scientific, MA, The US, #K1081).

### 4. Analytical methods

The growth of *Y. lipolytica* strains was monitored by measuring the OD_600_. Measured absorbance was converted into dry cell weight for *Y. lipolytica* as explained by Duman-Özdamar et al., 2022.

The total fatty acids were determined quantitatively with a gas chromatograph (GC), 7830B GC systems (Aligent, Santa Clara, CA, The US) equipped with a Supelco Nukol^TM^ 25357 column (30m x 530 µm x 1.0 µm; Sigma-Aldrich, St. Louis, MO, The US), hydrogen as a carrier gas. Samples were prepared as described by (Duman-Özdamar et al., 2022). Chloroform was evaporated under nitrogen gas and the remaining lipid in the tubes was dissolved in hexane before GC analysis.

Glycerol, citric acid, erythritol, mannitol, and arabitol were determined via HPLC analysis (Waters Alliance e2695, Milford, MA) with an RSpak KC811 column (ID = 8 mm, length = 300 mm, Shodex, NY) with a guard column RSpak KC-G (ID=6 mm and the length = 50 mm, Shodex, NY). The column was operated at 60°C with 3 mM H_2_SO_4_ as the mobile phase and a flow rate of 0.1 mL/min for 20 min. Peaks for components were detected and quantified with a refraction index detector (2414 RI Detector, Waters, Milford, MA). Peak integration and other chromatographic calculations were performed using Empower 3 software (Waters, Milford, MA). Identification and quantification of the corresponding compounds were achieved via comparisons to the standard curves (Figure S1).

### 5. Regression model

All computational analysis was performed with R version 4.0.2 (R Core Team, 2020). The relationship between the responses (*Y*), factors (*x*), and selected two-factor interactions were expressed by fitting a linear regression: 𝑌 = 𝛽_𝑜_ + ∑ 𝛽*_i_* 𝑥_𝑖_ + ∑ 𝛽_𝑖𝑗_𝑥_𝑖_𝑥_𝑗_. *βo* represents the interception coefficient, *βi* is the linear coefficient and *β_ij_* is the interaction coefficient.The quality of the regression equations was assessed according to the coefficient of determination (R^2^). Statistical analysis of the model was performed using Analysis of Variance (ANOVA) and p < 0.05 was considered significant.

## Results

### 1. Design

In the Design step of the DBTL approach (Figure 1), we evaluated the metabolic capabilities, and selected genes affecting lipid accumulation in *Y. lipolytica* for subsequent modulation, and designed experiments for the characterization of newly built strains.

**Figure 1.**
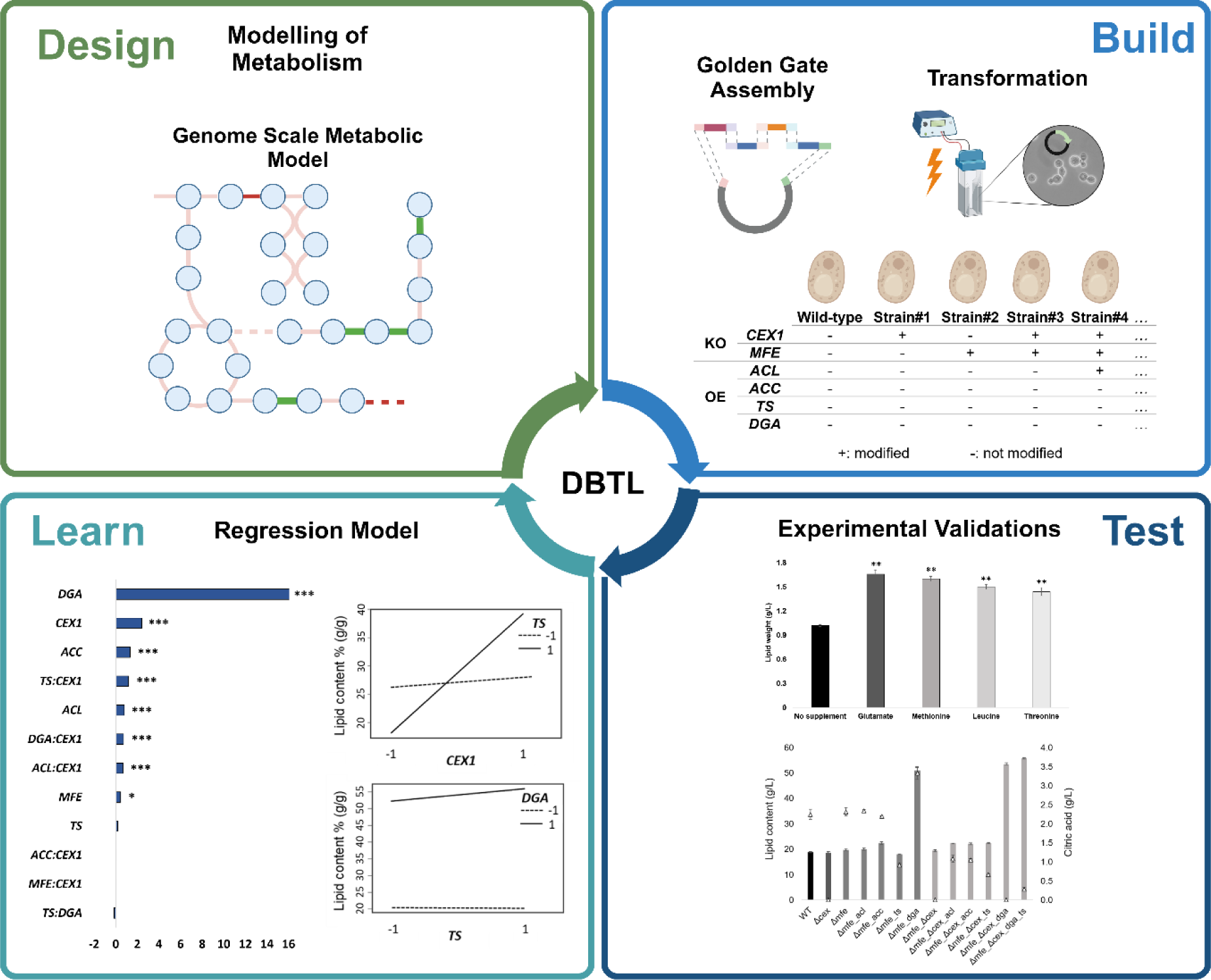
Overview of the DBTL approach **Design:** We analyzed the genome-scale metabolic model of *Y. lipolytica* (*i*Yali4, v4.1.2) to identify a genetic intervention strategy for enhancing the lipid content. **Build:** Plasmids were constructed via the golden gate assembly approach and confirmed via whole plasmid sequencing. Predicted target genes were overexpressed or knocked out via homologous recombination. **Test:** Predicted genetic interventions from the amino acid synthesis pathway were tested with wild-type (WT) by supplementing the corresponding amino acids to validate predictions and establish an efficient medium for increased lipid accumulation. Finally, the performance of the built transformants were characterized at C/N 140 minimal medium. **Learn:** Regression models were fitted to evaluate the main effects and two-factor interactions of genetic interventions on lipid content.

#### 1.1. Comparative Flux Sampling Analysis on GEM

Comparative Flux Sampling Analysis (CFSA) on the GEM model was performed to investigate genetic engineering strategies for improved lipid synthesis with *Y. lipolytica*. Flux distributions for each reaction were evaluated by simulating maximum production, and maximum growth (van Rosmalen *et al*., 2024). As a result, CFSA highlighted 70 overexpression targets (that could be clustered in 35 groups), 19 knock-out candidates (belonging to 17 groups), and 21 knock-down targets (belonging to 10 groups) (a complete list of targets and distribution plots are available at GitLab.

Reactions from the fatty acid synthesis and fatty acid elongation pathway, (fatty-acyl-CoA synthase (*ACS*), and stearoyl-CoA desaturase (*SCD*)) were suggested as overexpression targets (Table 1). Additionally, targets from pyruvate metabolism providing acetyl-CoA, (pyruvate kinase (*PK*), pyruvate decarboxylase (*PD*), acetaldehyde dehydrogenase (*ALDH*), and acetyl-CoA synthase (*ACS*)) and acetyl-CoA carboxylase (*ACC*), which converts acetyl-CoA to malonyl-CoA, were predicted as overexpression targets. Furthermore, several reactions from the phosphate pathway (PPP) (i.e., glucose 6-phosphate dehydrogenase (*G6PD*) and phosphogluconate dehydrogenase (*PGD*) providing NADPH, ribulose 5-phosphate 3-epimerase (*RPE1*) and transketolase (*TKL1, TKL2*)) were listed as overexpression targets. (Wasylenko et al., 2015) reported the oxidative PPP as a primary NADPH source for lipid synthesis and (Dobrowolski and Mirończuk, 2020) tested the increased NADPH availability via overexpression of *TKL1* and *DGA1* and reported around 1.5-fold increase in lipid content of *Y. lipolytica*.

**Table 1.**
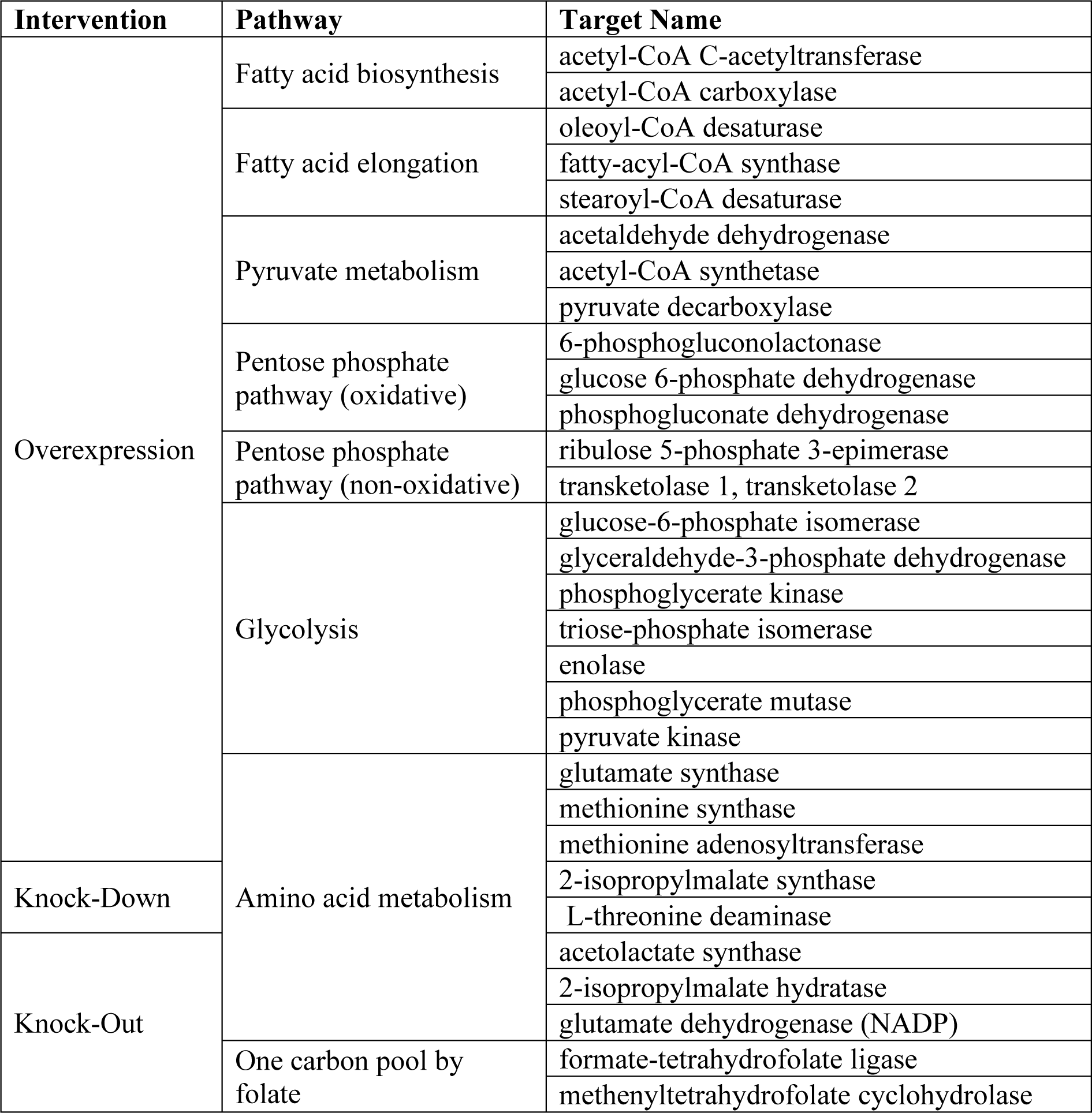
Selected genetic interventions (overexpression, knock down, knock-out) predicted to increase the lipid production of *Y. lipolytica* using CFSA on the genome-scale metabolic model. The complete list of predicted targets is available at GitLab.

CFSA predicted genetic intervention targets from amino acid metabolism including glutamate synthase (*GltS*), and methionine synthase (*MS*) for overexpression (Table 1). In addition, knock-out (acetolactate synthase (*ALS*), 2-isopropylmalate hydratase (*IPMS*)) and knock-down targets (2-isopropylmalate synthase (*LeuA*), L-threonine deaminase (*TD*)) related to threonine and leucine metabolism were highlighted. Lastly, reactions involved in one-carbon/methionine metabolism, methenyltetrahydrofolate cyclohydrolase (*MTHFC*), and formate-tetrahydrofolate ligase (*FTHFL*) were predicted as knock-out targets.

Subsequently, the predictions from the metabolic model intertwined with previous knowledge obtained for *Y. lipolytica* and *C. oleaginosus* (Duman-Özdamar et al., 2024). Combining all the outcomes led to the identification of amino acid supplements (glutamate, methionine, threonine, leucine), and four overexpression targets (*ACL1, ACC, TS* of *C. oleaginous,* and *DGA1* of *Y. lipolytica*) (Figure 2). CFSA or analysis of GEMs in general will not predict targets from competing pathways as their activity is not considered under the optimality conditions the model is set to operate. Therefore we additionally investigated the metabolism to investigate competing mechanisms such as the β-oxidation pathway and citrate secretion to the extracellular environment (Blazeck *et al*., 2013; Madzak, 2021). Selected overexpression targets were combined with the knock-out of citrate exporter protein (*CEX1*) (Odoni et al., 2019; Erian et al., 2020), and knock-out of a multifunctional enzyme (*MFE1*) catalyzing the second step of the β-oxidation pathway (Liu et al., 2021).

**Figure 2.**
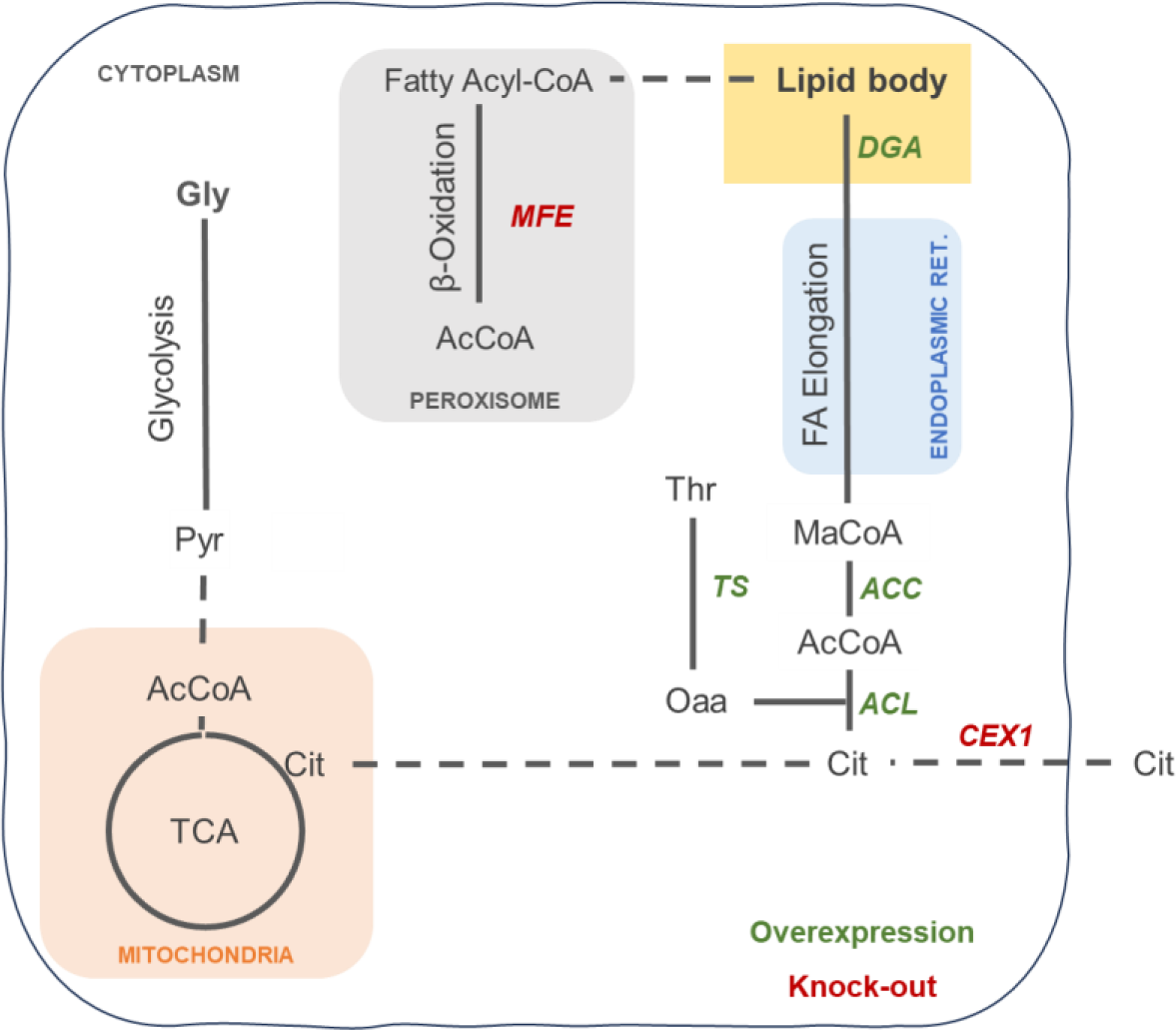
Summary of selected genetic interventions for improved lipid production in *Y. lipolytica*. TCA: tricarboxylic acid cycle, FA: fatty acids; Metabolite abbreviations: Gly: glycerol; Pyr: pyruvate, AcCoA: acetyl-coenzyme A, MaCoA: malonyl-coenzyme A, Cit: Citrate, Oaa: oxaloacetate, Thr: threonine.

### 2. Build

In the Build step, selected, genetic interventions were implemented (Figure 1). Plasmids were assembled using Golden Gate Assembly and confirmed via whole plasmid sequencing. Overexpression of *ACL, ACC, TS* (from *C. oleaginosus*), and *DGA1* (from *Y. lipolytica*) was achieved by knocking out *CEX1,* and/or *MFE1* via homologous recombination. In total, we constructed 12 strains in which we knocked out *CEX1, MFE1* individually and in combination (Figure 3A). After characterizing the background strains, we overexpressed the selected targets on *Δmfe* and *Δmfe_Δcex* background strains (Figure 3B-3C). Integration of the genes was confirmed via colony PCR (Figure S2).

**Figure 3.**
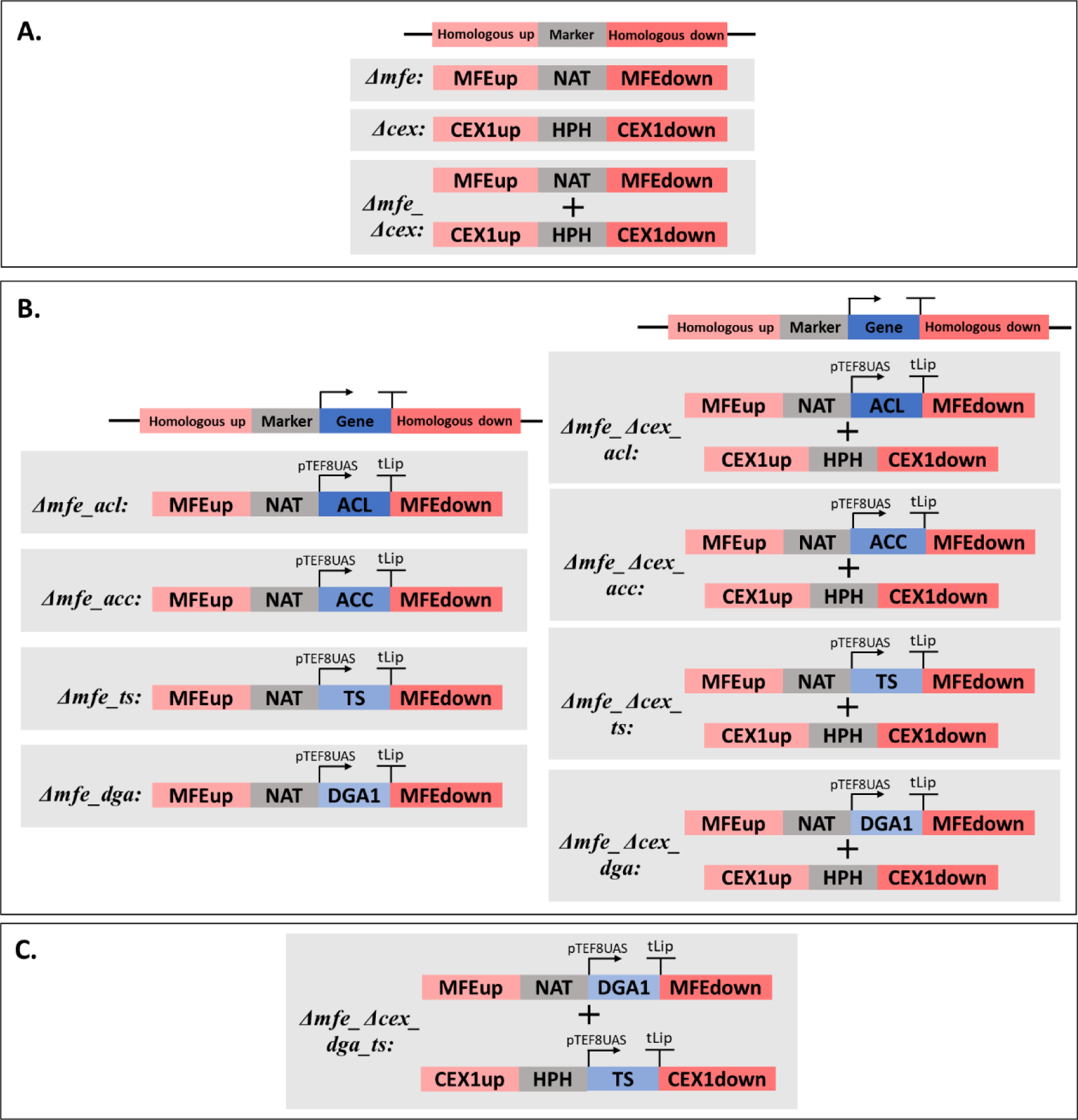
Overview of built step. **A) Build 1:** Background strains, *Δcex, Δmfe,* and *Δmfe_Δcex*, were built by transforming the assembled plasmids. **B) Build 2:** Single overexpression strains were built by overexpressing the selected targets (*ACL, ACC, TS, DGA1*) in *Δmfe* and *Δmfe_Δcex* background strains. **C) Build 3:** Double overexpression strain was built by overexpression of *DGA1* and *TS* in *Δmfe_Δcex*.

**Figure 3.**
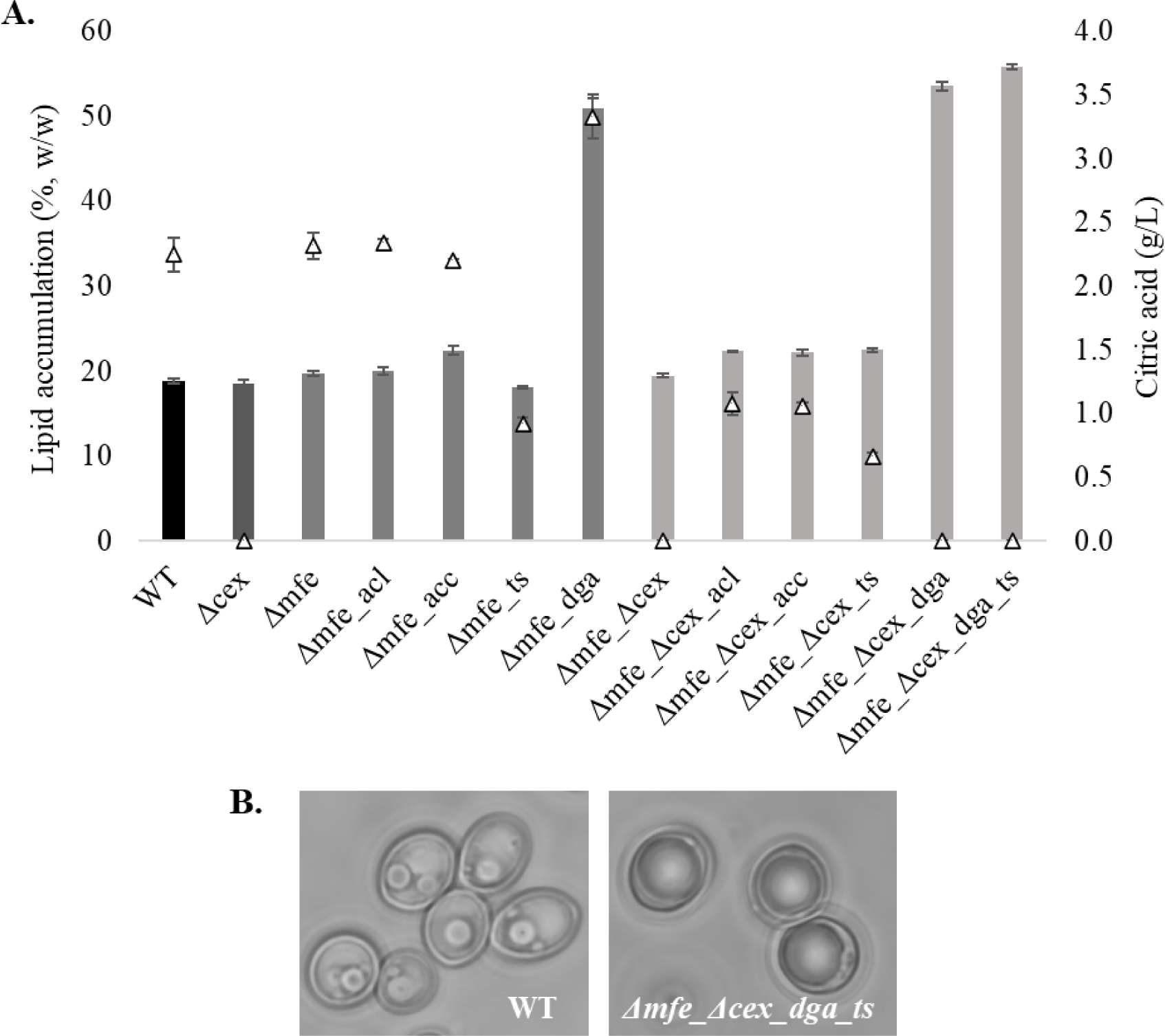
**A)** Lipid accumulation % (w/w) (bars), and extracellular citric acid (g/L) (triangles) of WT and *Y. lipolytica* transformants at C/N140 minimal medium at 120 hours. **B)** Microscope images of WT and *Δmfe_Δcex_dga_ts* at 120h showing that *Δmfe_Δcex_dga_ts* visibly more saturated with lipids.

### 3. Test

In the test step, we validated predictions from the amino acid synthesis pathway obtained on the design step via supplementation and observed the effect on the performance of *Y. lipolytica.* Furthermore constructed transformants in the build step were characterized at C/N140 cultivation medium containing glycerol as a carbon source and urea as a nitrogen source (Figure 1).

#### 3.1. Testing amino acid supplements

The predictions from the metabolic model related to the amino acid metabolism were experimentally validated by supplementing methionine, threonine, leucine, and glutamate into a nitrogen-limited cultivation medium (C/N140 g/g) (Duman-Özdamar et al., 2022). While these additions did not alter the lipid accumulation, total lipid production was 1.5-fold increased on average due to higher biomass concentrations (Table 2). Furthermore, biomass yield and lipid yield on glycerol were improved by around 1.6-fold with amino acid supplements. The addition of glutamate and threonine reduced extracellular citric acid by around 1.5-fold while it decreased by 1.6-fold with methionine and 2-fold with the addition of leucine. Supplementing amino acids did not affect the total content of saturated and unsaturated fatty acids, however, we observed around 7.5% shift of polyunsaturated fatty acids (PUFAs) to monounsaturated fatty acids (MUFAs) (Table S3).

**Table 2.** Lipid content, dry cell weight, biomass, and lipid yield on consumed glycerol of *Y. lipolytica,* and extracellular citrate concentrations upon supplementation of glutamate, methionine, leucine, or threonine into C/N140 (g/g) minimal medium (at 120h).

| Supplement | Dry cell weight (g/L) | Lipid content (% w/w) | $Y_{X/S}$ (g dcw/ g consumed glycerol) | $Y_{P/S}$ (g lipid/ g consumed glycerol) | Citric acid (g/L) |
| --- | --- | --- | --- | --- | --- |
| No supplement | $5.38 \pm 0.14$ | $18.94 \pm 0.34$ | $0.14 \pm 0.004$ | $0.026 \pm 0.001$ | $2.41 \pm 0.13$ |
| Glutamate | $9.34 \pm 0.55$ * | $17.85 \pm 0.66$ | $0.23 \pm 0.004$ * | $0.042 \pm 0.001$ * | $0.92 \pm 0.02$ * |
| Methionine | $8.80 \pm 0.21$ * | $18.20 \pm 0.50$ | $0.22 \pm 0.011$ * | $0.041 \pm 0.001$ * | $1.49 \pm 0.02$ * |
| Leucine | $8.24 \pm 0.29$ * | $18.18 \pm 0.32$ | $0.21 \pm 0.006$ * | $0.038 \pm 0.001$ * | $1.13 \pm 0.04$ * |
| Threonine | $7.74 \pm 0.23$ * | $18.63 \pm 0.68$ | $0.21 \pm 0.004$ * | $0.039 \pm 0.002$ * | $0.94 \pm 0.03$ * |
| Differences with control (no supplement) were evaluated using a t-test *: $p \leq 0.05$ | | | | | |

#### 3.2. Characterization of constructed *Y. lipolytica* transformants

Background strains (*Δcex*, *Δmfe,* and *Δmfe_Δcex*) and the overexpression of *ACL1, ACC, TS,* and *DGA* with *Δmfe* and *Δmfe_Δcex* backgrounds were characterized at C/N140 medium, which was identified as optimum C/N ratio for WT, in order to evaluate the effect of genetic interventions on lipid content, growth and extracellular citric acid concentration (Figure 3A) (Duman-Özdamar et al., 2022).

Secretion of citrate was blocked successfully with the knock-out of *CEX1,* while there was no significant effect on lipid content and dry cell weight (Table S4). We obtained a slight increase in lipid accumulation in *Δmfe* and *Δmfe_Δcex* compared to WT.

Overexpression of *ACL* resulted in a lipid content of 22.4% (w/w) when with simultaneous knock-out of *MFE1* and *CEX1*, which represents a 15% increase compared to *Δmfe_Δcex*. While *Δmfe_acc* and *Δmfe_Δcex_acc* accumulated around 22% (w/w) lipids and provided around 14% increase with respect to WT, a reduction in the lipid content was observed with *Δmfe_ts* (18.1%, w/w). On the other hand, *Δmfe_Δcex_ts* enhanced the lipid accumulation by 15% (22.45%, w/w). Although there was no extracellular citrate measured with *Δmfe_Δcex*, we detected a leakage of citrate in *Δmfe_Δcex*_*acl*, *Δmfe_Δcex*_*acc*, and *Δmfe_Δcex*_*ts* experiments which was declined approximately 2-fold compared to WT, *Δmfe_acl*, *Δmfe_acc*, and *Δmfe_ts*. In addition to citrate, mannitol (for all strains), erythritol (for all strains excluding *Δmfe*_*acl*), and arabitol (only *DGA1* overexpressed strains) were detected at 120h (Table S5).

The overexpression of *DGA* provided a sharp increase in lipid production*, Δmfe_dga* produced 51% (w/w), *Δmfe_Δcex_dga* accumulated 53.5% (w/w) lipids. There was no extracellular citrate detected with *Δmfe_Δcex_dga*, while *Δmfe_dga* secreted 1.4-fold higher citrate compared to *Δmfe*. Lastly, we overexpressed the *TS* and *DGA1* in the *Δmfe_Δcex* strain by considering improved lipid content and declined extracellular citrate concentrations in *Δmfe_dga, Δmfe_Δcex_dga,* and *Δmfe_Δcex_ts* strains. The ultimate increase was obtained with *Δmfe_Δcex_dga_ts* accumulated 56% (w/w) lipids (Figure 3B), which is a 2.8-fold increase in lipid content and a 3-fold improvement in Y_P/S_ compared to WT.

Regarding the fatty acid composition of accumulated lipids, the knock-out of *MFE1* resulted in a lower content of very long-chain fatty acids (VLC-FAs) in combination with all overexpressed genes and knock-out of *CEX1* compared to WT (Table S6). When *ACC* and *TS* overexpressed in *Δmfe* and *Δmfe_Δcex,* the content of saturated fatty acids (C16:0, C18:0) decreased by around 8%, and the content of MUFAs (C16:1, C18:1) increased by around 10%. Furthermore, *Δmfe_Δcex_acl* overexpression produced 7% lower saturated fatty acids and 8% higher MUFAs. On the other hand, we obtained the highest content of saturated fatty acids (on average 33.5%) with *Δmfe_dga, Δmfe_Δcex_dga,* and *Δmfe_Δcex_dga_ts*, while the content of MUFAs was around 12% higher compared to WT.

### 4. Learn

In the Learn step, we evaluated the impact of the tested genetic interventions both as main effects and selected 2-factor interactions (2Fi, *MFE:CEX1, ACL:CEX1, ACC:CEX1, TS:CEX1, DGA:CEX1, TS:DGA*) on the lipid content by fitting a second-order polynomial equation (Figure 1). ANOVA was conducted to evaluate the statistical significance and suitability of the model and the quality of the model fit was assessed using the coefficient of determination (R^2^ = 99.89%), and the significance was confirmed via p-value: < 2.2e-16 (Table S7).

The results highlighted the significant and positive effect of *DGA1, ACL, ACC* overexpression, *CEX1*, and *MFE1* knock-out on lipid content (Figure 4A). Among the indicated main effects, the greatest impact was observed by the modification of *DGA1* followed by *CEX1.* The outputs of the regression analysis showed positive significant interactions of *TS:CEX1* nevertheless, the main effect of *TS* and 2Fi of *TS:DGA* is insignificant on lipid content (Figure 4B). Furthermore, the model demonstrated a positive significant effect of *ACL:CEX1* and *DGA:CEX1*.

**Figure 4.**
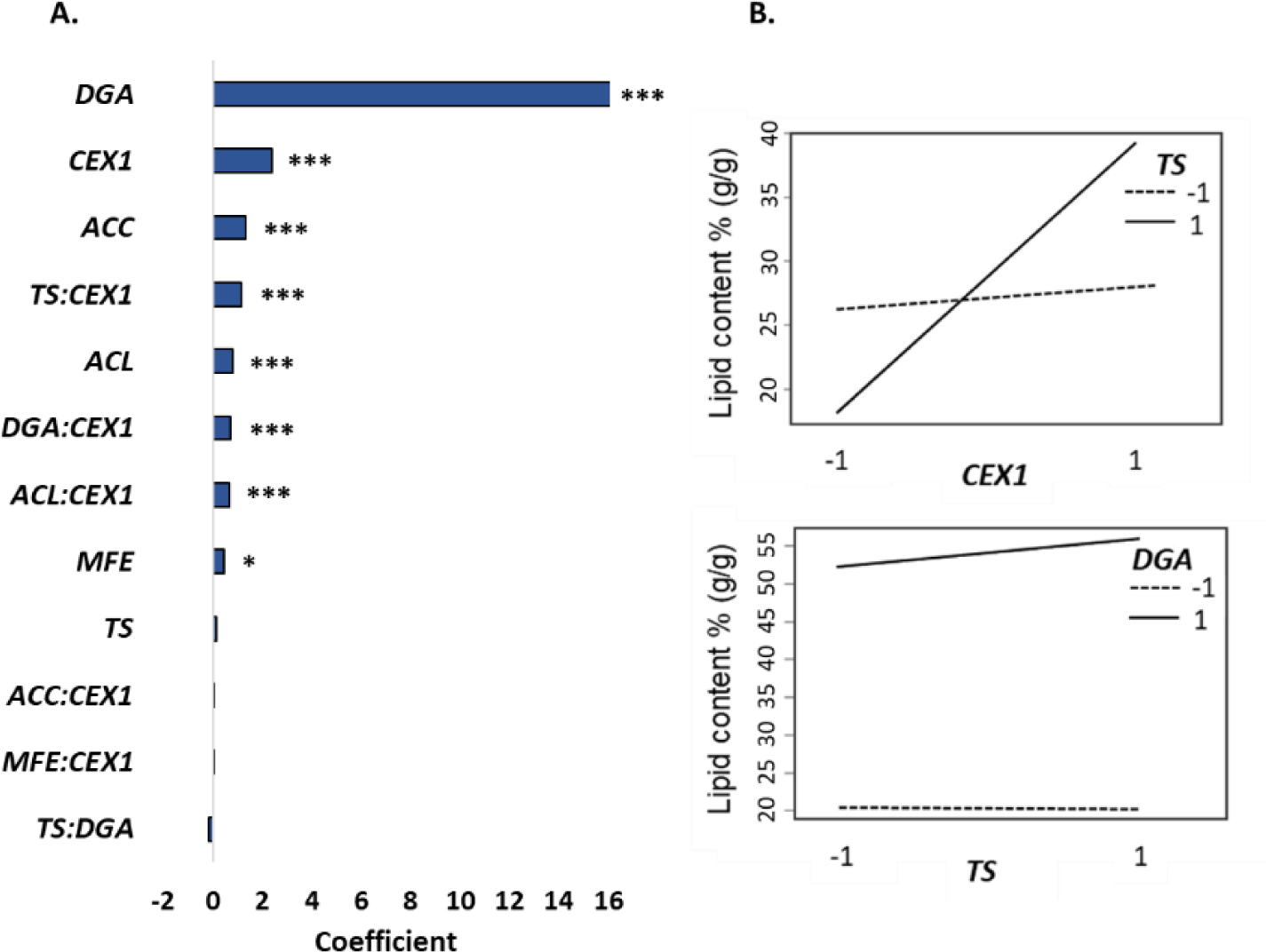
Regression model to evaluate the effects of tested genetic interventions and selected two-factor interactions on lipid content. A) Coefficients of the model. *** indicates corrected p-value ≤ 0.001, ** ≤ 0.01 and * ≤0.05. **B)** Interaction plot of *CEX1:TS* representing the positive significant interaction and interaction plot of *TS: DGA1* representing insignificant case.

## Discussion

An effort has been made to elucidate the lipid accumulation mechanism of *Y. lipolytica.* Developed genetic engineering tools, available GEM models and it is analysis for providing genetic engineering strategies enables a systematic approach to establish this oleaginous yeast as a sustainable fatty acid production platform (Beopoulos et al., 2009, 2011; Wang et al., 2020). In this study, we deployed the DBTL approach and focused on the lipid production potential of *Y. lipolytica* by intertwining the predictions from the GEM model, previous findings, and known bottlenecks for lipid accumulation of oleaginous yeasts with rounds of genetic interventions. We tested the effect of some amino acid supplements and characterized the built strains with shake flask experiments in N-limiting conditions. Statistical analysis was used to evaluate the gathered experimental data and evaluate the possible interactions between selected interventions on the lipid content of *Y. lipolytica.* The effect of the selected genetic interventions on the lipid content and the results from previous works were summarized in Table 3.

**Table 3.** Lipid content of engineered *Y. lipolytica*.

| Gene | Intervention | Gene source | Lipid content change | Carbon source | Reference |
| --- | --- | --- | --- | --- | --- |
| *CEX1* | Knock-out | - | No significant difference. | Glycerol | This work. |
| *MFE1* | Knock-out | - | Lipid content was increased from 16% to 19% (w/w). | Glucose | (Blazeck et al., 2013) |
|  | Knock-out | - | Lipid content was increased from 18.86% to 19.74% (w/w). | Glycerol | This work. |
| *ACL* | Knock-out | - | *Δacl1* mutant presents lower FA content and a higher citrate and mannitol production. | Glucose | (Dulermo et al., 2015) |
|  | Overexpression | *Mus musculus* | Lipid content increased from 7.3% to 23.1% (w/w). | Glycerol | (Zhang et al., 2014) |
|  |  | *Y. lipolytica* | No significant difference. | Glucose | (Blazeck et al., 2014) |
|  |  | *C. oleaginosus* | When combined with *Δmfe_Δcex*, lipid content was increased from 18.86% to 22.38% (w/w). | Glycerol | This work |
|  |  |  | When tested with *C. oleaginosus*, lipid content was increased from 37.8% to 51.8% (w/w). | Glycerol | (Duman-Özdamar et al., 2024) |
| *ACC* | Overexpression | *Y. lipolytica* | Lipid content was increased from 8.77% to 17.9% (w/w). | Glucose | (Tai and Stephanopoulos, 2013) |
|  |  |  | ACC1 and DGA1 overexpressed, Lipid content was increased from 8.77% to 41.4% (w/w). |  |  |
|  |  | *C. oleaginosus* | When combined with *Δmfe*, lipid content was increased from 18.86% to 22.39% (w/w). | Glycerol | This work |
|  |  | *C. oleaginosus* | When tested with *C. oleaginosus*, lipid content was increased from 37.8% to 48.9% (w/w). | Glycerol | (Duman-Özdamar et al., 2024) |
| *TS* | Supplement | - | Simulations on *iYL_2.0* resulted in a 2-fold increase in lipid productivity. | Glucose | (Wei et al., 2017) |
|  | Overexpression | *C. oleaginosus* | When combined with *Δmfe_Δcex*, lipid content was increased from 18.86% to 22.45% (w/w). | Glycerol | This work |
|  |  |  | When tested with *C. oleaginosus*, lipid content was increased from 37.8% to 51.4% (w/w). | Glycerol | (Duman-Özdamar et al., 2024) |
| Threonine synthesis pathway | Overexpression | *Y. lipolytica* | Overexpression of the threonine synthesis pathway decreased lipid content from 19% to 14.4% (w/w). | Glucose | (Park et al., 2020) |
| *DGA1* | Overexpression | *Y. lipolytica* | Around a 2-fold increase in lipid content (50% w/w). | Glucose | (Friedlander et al., 2016) |
|  |  |  | Lipid content was increased from 8.77% to 33.8% (w/w). | Glucose | (Tai and Stephanopoulos, 2013) |
|  |  |  | When combined with *Δmfe_Δcex*, lipid content was increased from 18.86% to 53.53% (w/w). | Glycerol | This work |
| *DGA1 + TS* | Overexpression | *Y. lipolytica* | When combined with *Δmfe_Δcex*, lipid content was increased from 18.86% to 55.87% (w/w). | Glycerol | This work |

When designing an efficient metabolic engineering strategy, it is important to understand not only the key reactions that are contributing to production but also the competing metabolic pathways that can affect productivity (Ratledge and Wynn, 2002; Beopoulos et al., 2009; Wen and Al Makishah, 2022). Blazeck et al. (2013) tested the deletion of *MFE1* and *PEX10* related to the β-oxidation pathway and reported that the lipid accumulation increased by 19% (Table 3). Additionally, secretion of citric acid to the extracellular environment is identified as one of the main limitations on the utilization of intracellular citrate for lipid biosynthesis especially when *Y. lipolytica* grows on glycerol (Moeller et al., 2011; Sagnak et al., 2018; Wang et al., 2020). Recently, the first citrate exporter of *Y. lipolytica*, was identified (Erian et al., 2020). In our metabolic engineering strategy, we implemented the knock-out of *MFE1* and *CEX1* to increase the intracellular citrate availability and prevent cleavage of fatty acid via the β-oxidation pathway. In the background strains *Δmfe and Δmfe_ Δcex*, a slight increase in lipid content was observed, and we were able to block citrate secretion in the *Δcex, Δmfe_ Δcex knock-out strains*.

Analysis of the metabolic model predicted *ACC* as a suitable target, which initiates lipid synthesis by providing malonyl-CoA. Tai and Stephanopoulos (2013) reported that overexpression of *ACC* improved lipid content by around 2-fold. Our previous work showed that *ACC* overexpression in *C. oleaginosus* increased lipid content by 30% (Table 4) (Duman-Özdamar et al., 2024). In this work, we observed that lipid accumulation of *Δmfe_acc* improved by 19% and *Δmfe_Δcex_acc* by 18%. Furthermore, the fitted polynomial model represented a positive significant effect of *ACC* on lipid content but no significant interaction between ACC:CEX1. This suggests that preventing the secretion of citrate is not beneficial to increasing accumulated lipids when overexpressing ACC.

In addition to the ACC, ACL was identified as a critical reaction. Dulermo et al. (2015) reported that the knock-out of *ACL* in *Y. lipolytica* led to repression of the lipid synthesis. On the other hand, overexpressing the *ACL* of *Y. lipolytica* did not alter the lipid synthesis that was encountered by overexpressing the *ACL* of *Mus musculus* (Blazeck et al., 2014; Zhang et al., 2014). This was explained by the lower citrate affinity of *Y. lipolytica ACL* compared to the *Mus musculus ACL*. Also, the authors reported that this overexpression redirected a significant amount of the cytosolic citrate to the lipid synthesis pathway. As the *ACL* of *Y. lipolytica* showed lower affinity to citrate, we overexpressed *ACL* of *C. oleaginous* providing a 37% increase in the lipid content of *C. oleaginosus* (Duman-Özdamar et al., 2024). We obtained a significant increase in lipid accumulation with only *Δmfe_Δcex_acl* (by 19%), while there was around 2-fold declined extracellular citric acid compared to WT. Statistical analysis indicated a positive significant interaction of CEX:ACL on lipid content that shows the simultaneous modification of their expression levels is a successful approach to increased lipid synthesis.

In addition to predictions from the fatty acid synthesis pathway, CFSA predicted genetic intervention targets from the amino acid synthesis pathway related to glutamate, methionine, threonine, and leucine metabolism. Wei et al. (2017) simulated the supplement of L-serine, L-threonine, and L-aspartate and reported the supplements enhanced TAG production. We tested glutamate, methionine, leucine, and threonine as medium supplements and observed that the amino acid supplements enhanced the total lipid (g/L) around 1.6-fold however, it did not alter the lipid content. Furthermore, Kim et al. (2019) performed simulations on the GEM model of *Y. lipolytica* and reported that the threonine synthesis pathway was predicted as an overexpression target for improved lipid content. On the other hand, Park et al. (2020) indicated that the lipid content of *Y. lipolytica* declined due to the overexpression of homoserine kinase, and threonine synthase (Table 3). In our previous work, we overexpressed *TS* in *C.oleaginosus* and obtained a 36% increase in lipid content (Duman-Özdamar et al., 2024). Therefore the effect of *TS* overexpression was tested for *Y. lipolytica* by constructing *Δmfe_ts* and *Δmfe_Δcex_ts* transformants. While the lipid content of *Δmfe_ts* declined by around 5%, we obtained around 19% increase in the lipid content of *Δmfe_Δcex_ts*. In both cases measured extracellular citrate concentrations declined by 2.4-fold with *Δmfe_ts* and 3.4-fold with *Δmfe_Δcex_ts.* Besides these results, our model represented that the main effect of *TS* on lipid content is not significant however interaction of TS:CEX1 has a positive effect on lipid content. In all, these results indicate when the intracellular citrate concentration is higher, *TS* overexpression supports lipid accumulation possibly by balancing the over-production of oxaloacetate (Kim et al., 2019; Duman-Özdamar et al., 2024).

No extracellular citric acid was found in the *Δcex* strain. On the other hand, when *ACL, ACC,* and *TS* were overexpressed we detected secreted citric acid (up to 1 g/L, 2-fold lower than WT). Colony PCR was performed at the end of cultivation for all *Δcex* strains and we again confirmed the *Δcex* genotype. Altogether this indicates other exporters are able to transport citrate in the genome of this yeast (Lazar et al., 2017; Erian et al., 2020).

Another selected gene was *DGA1*, catalyzing the last step of TAG biosynthesis of *Y. lipolytica* (Beopoulos et al., 2012; Tai and Stephanopoulos, 2013)*. DGA1* overexpression showed a push effect of fatty acid synthesis and provided around a 2-fold increase in lipid accumulation (Tai and Stephanopoulos, 2013; Friedlander et al., 2016). In our work, we overexpressed *DGA1* in *Δmfe* and *Δmfe_Δcex* which resulted in a 2.7-fold and 2.8-fold increase in lipid content respectively. *DGA1* has the highest positive effect on lipid content among tested genetic interventions and DGA: CEX1 represented a positive interaction. Ultimately we combined the overexpression of *TS,* as it was leading to the lowest extracellular citrate concentrations compared to other targets, and *DGA1* in the *Δmfe_Δcex* background which resulted in around 200% increase in lipid content (56% w/w) and a 230% increase in Y_P/S_ (0.085 g/g).

A vast amount of work has been done in glucose-based medium with *Y. lipolytica* (Gajdoš et al., 2017; Wang et al., 2020). These results represent the potential of *Y. lipolytica* in the glycerol-based medium. Altogether, we believe the strain developed and the findings in this study have remarkable potential, especially for the conversion of glycerol-containing side streams i.e. crude glycerol derived from biodiesel production, fat saponification due to producing lipids with higher yields (André et al., 2009; Dobrowolski et al., 2016; Tsirigka et al., 2023).

## Conclusion

In this study, we demonstrated a comprehensive and systematic approach to enhance lipid production in *Y. lipolytica* using the Design-Build-Test-Learn (DBTL) methodology. By integrating genetic intervention predictions from the genome-scale metabolic model (GEM) with experimental validations and finally fitting a second-order polynomial model, we achieved significant improvements in lipid accumulation. Our work highlighted the crucial roles of *ACC, ACL, TS,* and *DGA1* and the interaction of these genes with increased intracellular citrate availability in lipid biosynthesis. Furthermore, we observed the positive effects of amino acid supplementation on total lipid and lipid yield on glycerol. Notably, overexpression of *DGA1* in *Δmfe* and *Δmfe_Δcex* backgrounds led to a remarkable 2.7-fold and 2.8-fold increase in lipid content, respectively, while the combination of *TS* and *DGA1* overexpression in *Δmfe_Δcex* background resulted in 200% increase in lipid content and a 230% increase in Y_P/S_. These results underscore the potential of *Y. lipolytica* as a sustainable alternative for fatty acid production. The insights gained from this study not only advance our understanding of lipid metabolism in oleaginous yeasts but also pave the way for industrial applications, particularly in utilizing glycerol-containing by-products for bio-based lipid production. Our findings contribute to the ongoing efforts to develop environmentally friendly and economically viable microbial oil production platforms, addressing the pressing need for sustainable alternatives to palm oil.

## Data availability statement

Supplementary files related to CFSA are available at <u>GitLab</u>., https://gitlab.com/wurssb/Modelling/sampling-tools, supplementary figures and tables are available at the end of the file.

## Author contributions

All authors conceived and designed the study. ZEDÖ and MSD performed the data analysis. ZEDÖ drafted the manuscript and performed the experiments. VAPMdS, JH, and MSD acquired project funding, and VAPMdS, JH, MSD, and MKJ conceived and supervised the research. All authors reviewed and edited the study. All authors read and approved the final manuscript.

## Acknowledgments

We thank Faezah Feyzi for her valuable contribution to the selection of the genome-scale metabolic model and the medium supplement experiments. Graphical abstract and Figure 1 were created with BioRender.com.

## Supplementary Material

**Figure S1.**
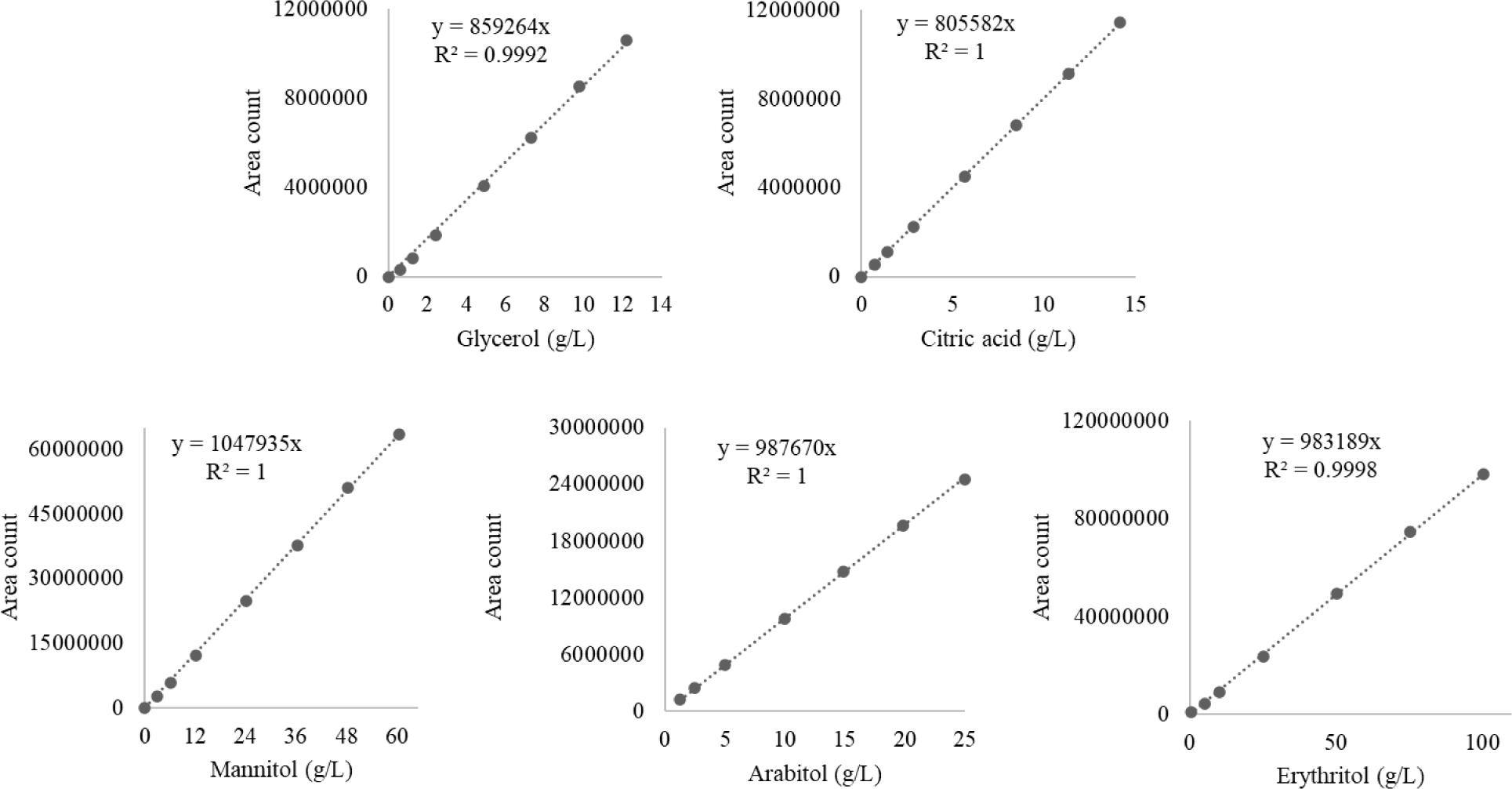
Calibration curve of glycerol, citric acid, mannitol, arabitol, and erthritol for calculating their concentrations in the medium.

**Figure S2.**
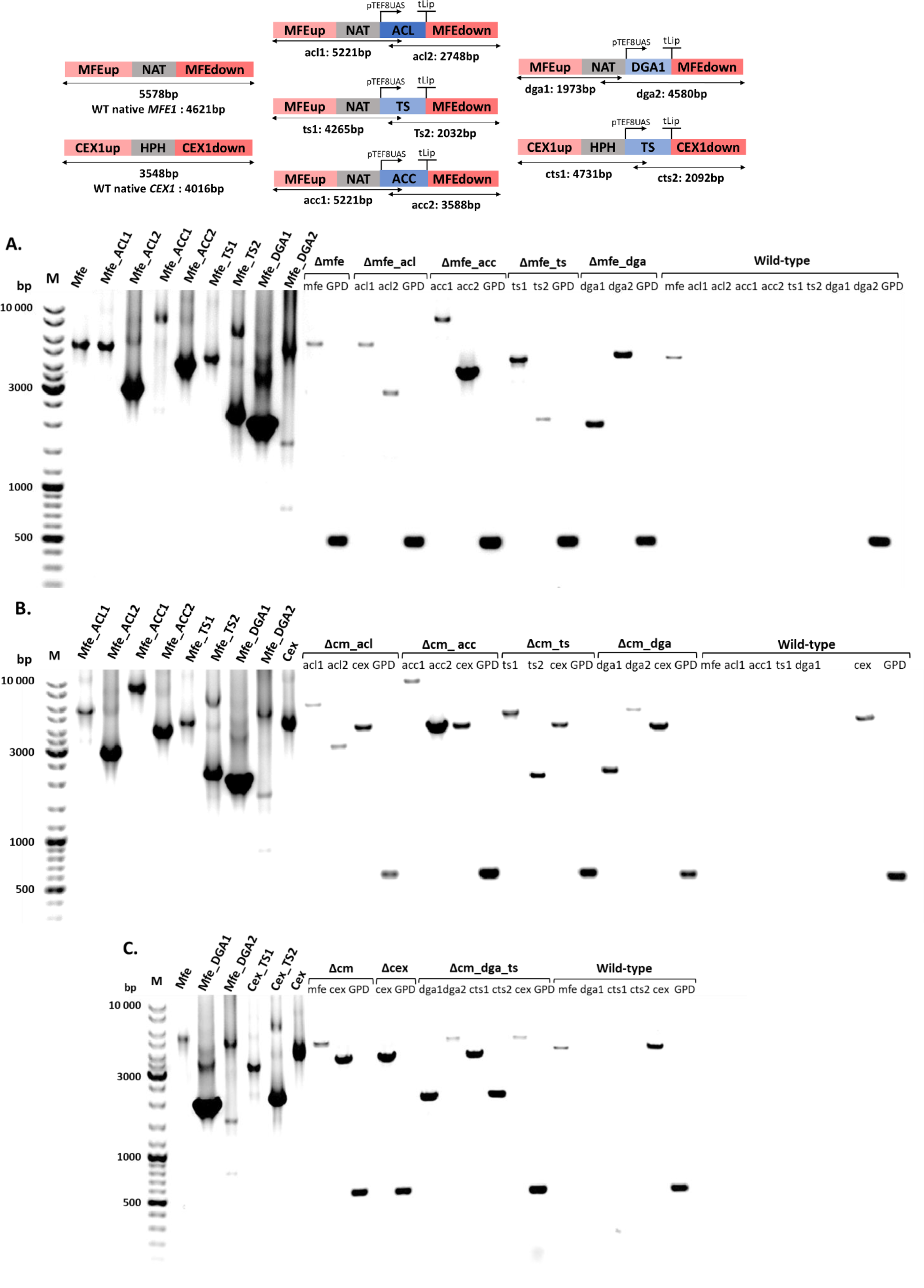
Colony PCR products were run on 1 % agarose gel. Positive controls were prepared by using corresponding primers and plasmids. Quality of gDNA in PCR reaction was assessed by using primers specific for the glyceraldehyde-3-phosphate dehydrogenase gene (*GPD,* Accession: XM_500444). Expected lengths were indicated per construct. **A)** Agarose gel photo of *Δmfe* strains **B)** Agarose gel photo of *Δcex_Δmfe* strains **C)** Agarose gel photo of *Δcex_Δmfe_dga_ts*.

**Table S1.**
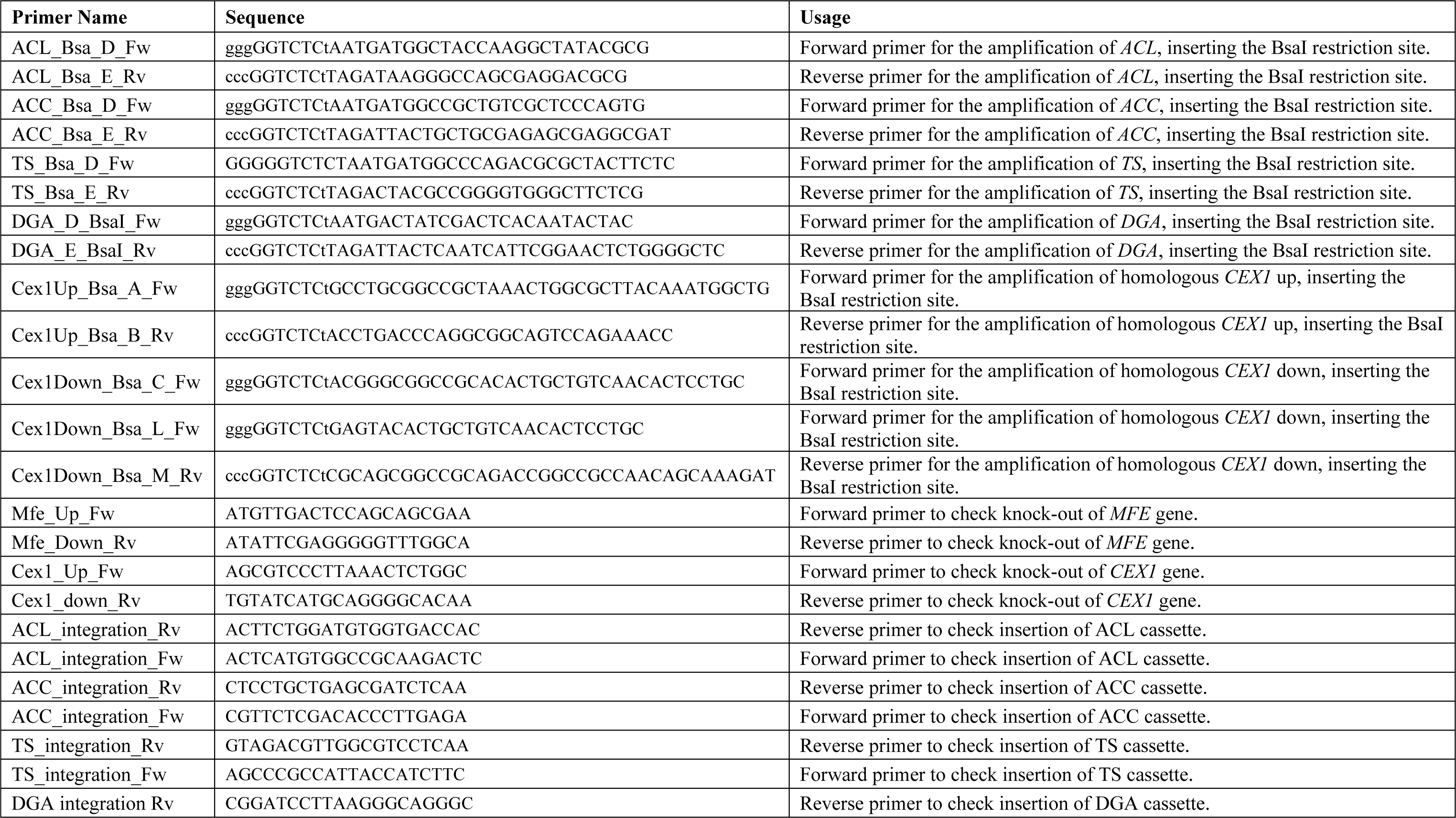
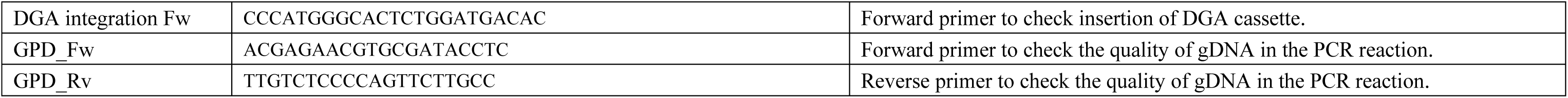
Primers designed for cPCR and PCR amplification of genes.

**Table S2.** Plasmids used to assamble the cassettes for knock-out and overexpress the genetic interventions and built strains.

| Strain | Plasmids used in assembly | Genotype/Assembled plasmids |
| --- | --- | --- |
| *Δmfe* | pSB1A3-GB3 + pCR4Blunt-TOPO-MFE-NotI_Up + pCR4Blunt-TOPO-M-Nat + pCR4Blunt-TOPO-MFE-NotI_Down | pSB1A3-MfeUP-NAT-MfeDOWN |
| *Δcex* | pSB1A3-GB3 + pCRBlunt-CEX1_Up + pCR4Blunt-TOPO-M-hph + pCRBlunt-CEX1_CM_Down | pSB1A3-Cex1UP-HPH-Cex1DOWN |
| *Δmfe_acl* | pSB1A3-GB3 + pCR4Blunt-TOPO-MFE-NotI_Up + pCR4Blunt-TOPO-M-Nat+ pCR4Blunt-TOPO- P1 TEF-8UAS + pCRBlunt-ACL + pCR4Blunt-TOPO-TLip2 (E-L) + pCR4Blunt-TOPO-MFE-NotI_Down | pSB1A3-MfeUP-NAT-pTEF8UAS-ACL-tLip2-MfeDOWN |
| *Δmfe_acc* | pSB1A3-GB3 + pCR4Blunt-TOPO-MFE-NotI_Up + pCR4Blunt-TOPO-M-Nat+ pCR4Blunt-TOPO- P1 TEF-8UAS + pCRBlunt-ACC + pCR4Blunt-TOPO-TLip2 (E-L) + pCR4Blunt-TOPO-MFE-NotI_Down | pSB1A3-MfeUP-NAT-pTEF8UAS-ACL-tLip2-MfeDOWN |
| *Δmfe_ts* | pSB1A3-GB3 + pCR4Blunt-TOPO-MFE-NotI_Up + pCR4Blunt-TOPO-M-Nat+ pCR4Blunt-TOPO- P1 TEF-8UAS + pCRBlunt-TS + pCR4Blunt-TOPO-TLip2 (E-L) + pCR4Blunt-TOPO-MFE-NotI_Down | pSB1A3-MfeUP-NAT-pTEF8UAS-ACL-tLip2-MfeDOWN |
| *Δmfe_dga* | pSB1A3-GB3 + pCR4Blunt-TOPO-MFE-NotI_Up + pCR4Blunt-TOPO-M-Nat+ pCR4Blunt-TOPO- P1 TEF-8UAS + pCRBlunt-DGA + pCR4Blunt-TOPO-TLip2 (E-L) + pCR4Blunt-TOPO-MFE-NotI_Down | pSB1A3-MfeUP-NAT-pTEF8UAS-ACL-tLip2-MfeDOWN |
| *Δmfe_ Δcex* | pSB1A3-GB3 + pCR4Blunt-TOPO-MFE-NotI_Up + pCR4Blunt-TOPO-M-Nat + pCR4Blunt-TOPO-MFE-NotI_Down | pSB1A3-MfeUP-NAT-MfeDOWN |
|  | pSB1A3-GB3 + pCRBlunt-CEX1_Up + pCR4Blunt-TOPO-M-hph + pCRBlunt-CEX1_CM_Down | pSB1A3-Cex1UP-HPH-Cex1DOWN |
| *Δmfe_ Δcex_acl* | pSB1A3-GB3 + pCRBlunt-CEX1_Up + pCR4Blunt-TOPO-M-hph + pCRBlunt-CEX1_CM_Down | pSB1A3-Cex1UP-HPH-Cex1DOWN |
|  | pSB1A3-GB3 + pCR4Blunt-TOPO-MFE-NotI_Up + pCR4Blunt-TOPO-M-Nat+ pCR4Blunt-TOPO- P1 TEF-8UAS + pCRBlunt-ACL + pCR4Blunt-TOPO-TLip2 (E-L) + pCR4Blunt-TOPO-MFE-NotI_Down | pSB1A3-MfeUP-NAT-pTEF8UAS-ACL-tLip2-MfeDOWN |
| *Δmfe_ Δcex_acc* | pSB1A3-GB3 + pCRBlunt-CEX1_Up + pCR4Blunt-TOPO-M-hph + pCRBlunt-CEX1_CM_Down | pSB1A3-Cex1UP-HPH-Cex1DOWN |
|  | pSB1A3-GB3 + pCR4Blunt-TOPO-MFE-NotI_Up + pCR4Blunt-TOPO-M-Nat+ pCR4Blunt-TOPO- P1 TEF-8UAS + pCRBlunt-ACC + pCR4Blunt-TOPO-TLip2 (E-L) + pCR4Blunt-TOPO-MFE-NotI_Down | pSB1A3-MfeUP-NAT-pTEF8UAS-ACC-tLip2-MfeDOWN |
| *Δmfe_Δcex_ts* | pSB1A3-GB3 + pCRBlunt-CEX1_Up + pCR4Blunt-TOPO-M-hph + pCRBlunt-CEX1_CM_Down | pSB1A3-Cex1UP-HPH-Cex1DOWN |
|  | pSB1A3-GB3 + pCR4Blunt-TOPO-MFE-NotI_Up + pCR4Blunt-TOPO-M-Nat+ pCR4Blunt-TOPO- P1 TEF-8UAS + pCRBlunt-TS + pCR4Blunt-TOPO-TLip2 (E-L) + pCR4Blunt-TOPO-MFE-NotI_Down | pSB1A3-MfeUP-NAT-pTEF8UAS-TS-tLip2-MfeDOWN |
| *Δmfe_Δcex_dga* | pSB1A3-GB3 + pCRBlunt-CEX1_Up + pCR4Blunt-TOPO-M-hph + pCRBlunt-CEX1_CM_Down | pSB1A3-Cex1UP-HPH-Cex1DOWN |
|  | pSB1A3-GB3 + pCR4Blunt-TOPO-MFE-NotI_Up + pCR4Blunt-TOPO-M-Nat+ pCR4Blunt-TOPO- P1 TEF-8UAS + pCRBlunt-DGA + pCR4Blunt-TOPO-TLip2 (E-L) + pCR4Blunt-TOPO-MFE-NotI_Down | pSB1A3-MfeUP-NAT-pTEF8UAS-DGA-tLip2-MfeDOWN |
| *Δmfe_Δcex_dga_ts* | pSB1A3-GB3 + pCRBlunt-CEX1_Up + pCR4Blunt-TOPO-M-hph + pCR4Blunt-TOPO- P1 TEF-8UAS + pCRBlunt-TS + pCR4Blunt-TOPO-TLip2 (E-L) + pCRBlunt-CEX1_LM_Down | pSB1A3-Cex1UP-HPH-pTEF8UAS-TS-tLip2-Cex1DOWN |
|  | pSB1A3-GB3 + pCR4Blunt-TOPO-MFE-NotI_Up + pCR4Blunt-TOPO-M-Nat+ pCR4Blunt-TOPO- P1 TEF-8UAS + pCRBlunt-DGA + pCR4Blunt-TOPO-TLip2 (E-L) + pCR4Blunt-TOPO-MFE-NotI_Down | pSB1A3-MfeUP-NAT-pTEF8UAS-DGA-tLip2-MfeDOWN |

**Table S3.** Fatty acid profile of *Y. lipolytica* grown at minimal medium with or without supplement (glutamate, methionine, leucine, threonine) at 120h. The fatty acid profile of palm oil was added for direct comparison. MUFAs: Monounsaturated fatty acids, PUFAs: Polyunsaturated fatty acids, VLC-FAs: Very long chain fatty acids.

| Fatty acid profile (%) |  |  |  |  |  |  |  |  |  |  |  |  |  |  |  |  |  |
| --- | --- | --- | --- | --- | --- | --- | --- | --- | --- | --- | --- | --- | --- | --- | --- | --- | --- |
| Supplement | C10:0 | C16:0 | C16:1 | C16:2 | C16:3 | C18:0 | C18:1 | C18:2 | C18:3 | C20:0 | C20:1 | C22:0 | C24:0 | Saturated FAs | MUFAs | PUFAs | VLC-FAs |
| No supplement | 0.35 ± 0.04 | 16.23 ± 0.90 | 9.26 ± 0.33 | 0.57 ± 0.02 | 0.70 ± 0.01 | 10.42 ± 0.55 | 37.42 ± 1.27 | 21.14 ± 0.03 | 0.29 ± 0.02 | 0.45 ± 0.01 | - | 0.59 ± 0.02 | 2.59 ± 0.14 | 30.63 ± 1.65 | 46.67 ± 1.61 | 22.70 ± 0.06 | 3.18 ± 0.16 |
| Glutamate | 0.41 ± 0.05 | 14.45 ± 0.52 | 10.45 ± 0.61 | - | 1.15 ± 0.10 | 11.52 ± 1.00 | 42.92 ± 0.81 | 16.25 ± 1.23 | - | - | - | 0.50 ± 0.06 | 2.34 ± 0.24 | 29.22 ± 1.79 | 53.37 ± 1.31 ** | 17.41 ± 1.17 ** | 2.84 ± 0.30 |
| Methionine | 0.53 ± 0.02 | 15.71 ± 0.20 | 9.68 ± 0.46 | - | 1.26 ± 0.11 | 10.82 ± 0.33 | 44.14 ± 0.69 | 14.91 ± 0.43 | - | - | - | 0.64 ± 0.08 | 2.32 ± 0.12 | 30.02 ± 0.71 | 53.82 ± 1.12 ** | 16.17 ± 0.54 ** | 2.96 ± 0.19 |
| Leucine | 0.55 ± 0.12 | 14.86 ± 0.53 | 8.60 ± 0.50 | - | 1.09 ± 0.11 | 13.97 ± 0.89 | 45.64 ± 0.71 | 11.02 ± 0.49 | - | 0.73 ± 0.07 | - | 0.61 ± 0.05 | 2.93 ± 0.08 | 33.65 ± 1.32 | 54.24 ± 0.95 ** | 12.11 ± 0.56 ** | 3.53 ± 0.12 |
| Threonine | 0.60 ± 0.17 | 15.10 ± 0.39 | 10.14 ± 0.54 | - | 1.14 ± 0.02 | 12.20 ± 0.47 | 44.26 ± 0.50 | 13.31 ± 0.79 | - | - | - | 0.58 ± 0.03 | 2.65 ± 0.16 | 31.14 ± 0.48 | 54.41 ± 0.29 ** | 14.45 ± 0.77 ** | 3.23 ± 0.18 |
| Palm (range) (Montoya et al., 2013) | - | 32.6–46.0 | 0.0–0.5 | - | - | 1.7–7.6 | 35.1–53.4 | 7.0–15.0 | 0.2–0.6 | 0.1–0.5 | 0.1–0.2 | – | – | 36.7–51.5 | 35.3–53.8 | 7.3–15.5 | - |
| Differences with control (no supplement) were evaluated using a t-test *: $p \leq 0.05$, **: $p \leq 0.01$. | | | | | | | | | | | | | | | | | |

**Table S4.**
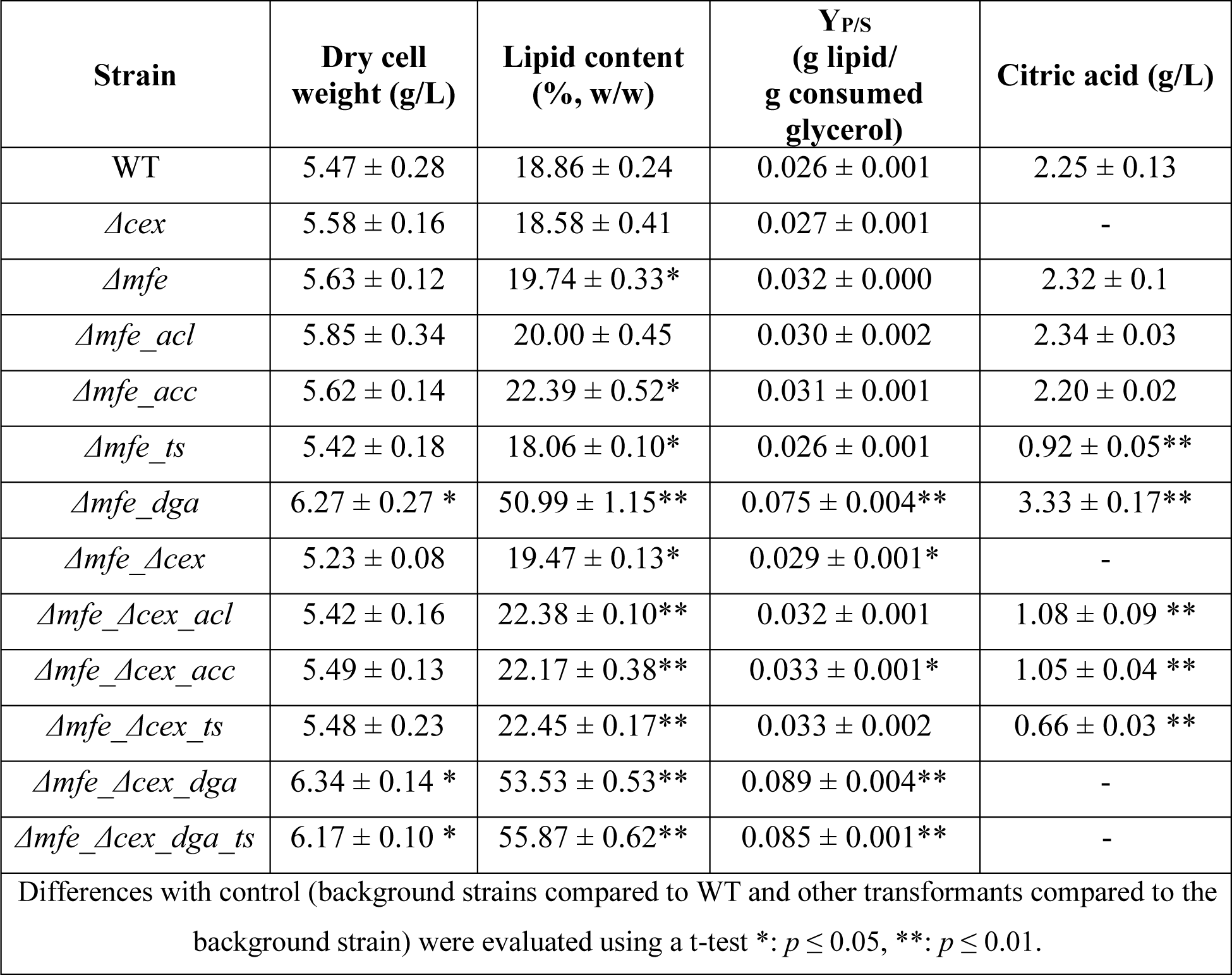
Lipid content, dry cell weight, lipid weight, biomass, and lipid yield on consumed glycerol of built *Y. lipolytica* strains and extracellular citrate concentrations at C/N140 (g/g) minimal medium (at 120h).

**Table S5.**
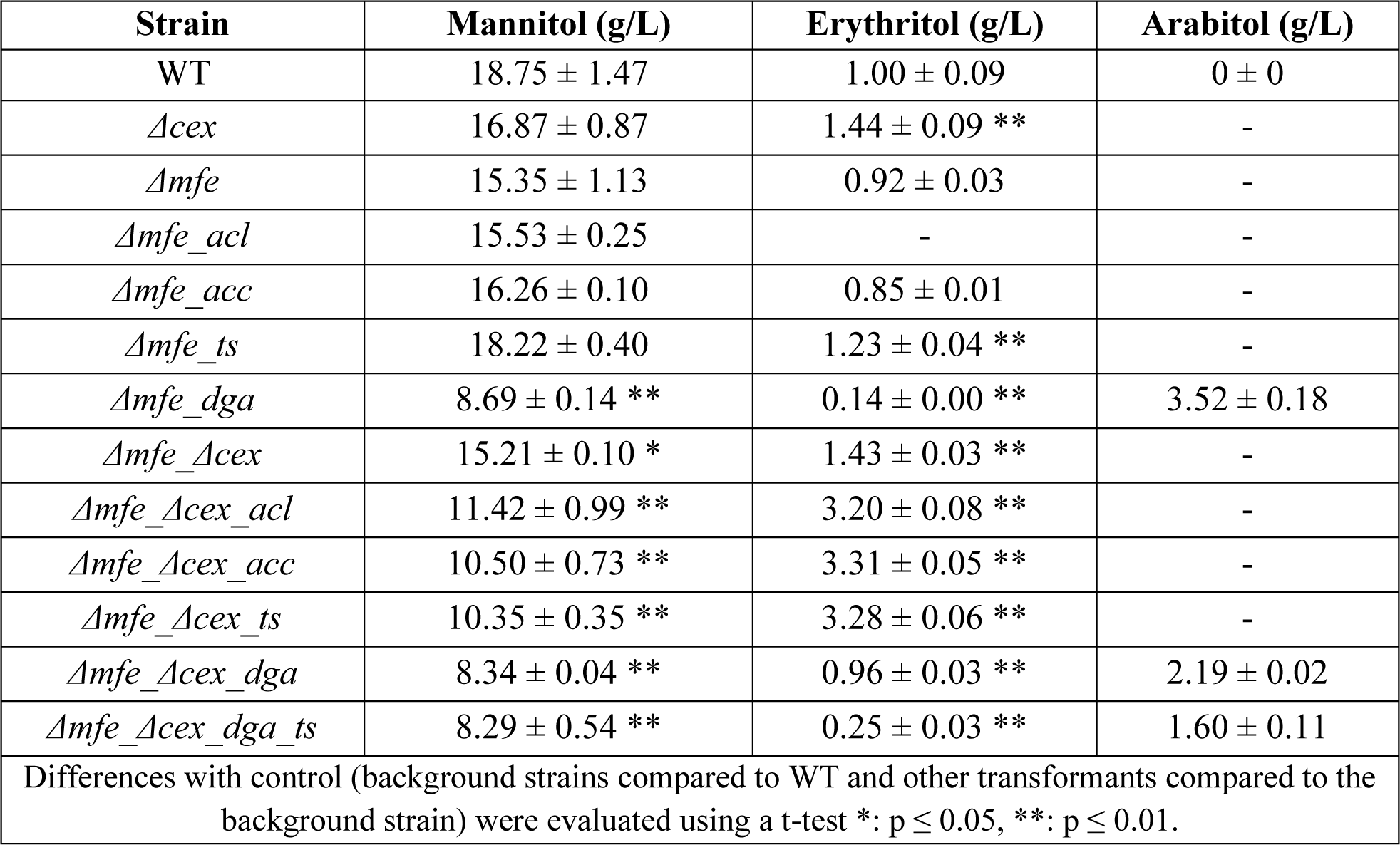
Measured erythritol and arabitol concentration of *Y. lipolytica* strains and at C/N140 (g/g) minimal medium (at 120h).

**Table S6.** Fatty acid profile of transformants and WT grown at C/N140 at 120 h. The fatty acid profile of palm oil was added for direct comparison. MUFAs: Monounsaturated fatty acids, PUFAs: Polyunsaturated fatty acids. Differences with control (WT) were evaluated using a t-test *: *p* ≤ 0.05, **: *p* ≤ 0.01.

| Strain | Fatty Acid Profile (%) |  |  |  |  |  |  |  |  |  |  |  |  |  |  |  |  |  |  |
| --- | --- | --- | --- | --- | --- | --- | --- | --- | --- | --- | --- | --- | --- | --- | --- | --- | --- | --- | --- |
|  | C10:0 | C14:0 | C16:0 | C16:1 | C16:2 | C16:3 | C18:0 | C18:1 | C18:2 | C18:3 | C20:0 | C20:1 | C22:0 | C22:1 | C24:0 | Saturated FA | MUFAs | PUFAs | VLC-FAs |
| WT | 0.35 ± 0.01 | 0.27 ± 0.02 | 16.63 ± 0.31 | 9.31 ± 0.32 | 0.55 ± 0.04 | 0.75 ± 0.02 | 10.86 ± 0.29 | 36.34 ± 0.22 | 20.75 ± 0.37 | 0.24 ± 0.01 | 0.42 ± 0.01 | 0.85 ± 0.15 | 0.56 ± 0.01 | - | 2.13 ± 0.15 | 31.21 ± 0.53 | 46.50 ± 0.32 | 22.29 ± 0.34 | 2.68 ± 0.14 |
| *Δcex* | 0.42 ± 0.05 | 0.34 ± 0.05 | 16.49 ± 1.17 | 10.34 ± 0.33 | 0.45 ± 0.01 | 0.69 ± 0.02 | 8.39 ± 0.48 | 39.21 ± 0.20 | 21.12 ± 0.20 | - | 0.53 ± 0.01 | 0.65 ± 0.06 | 0.40 ± 0.01 | - | 2.29 ± 0.21 | 28.86 ± 0.62 * | 50.20 ± 0.49 ** | 22.34 ± 0.62 | 2.69 ± 0.20 |
| *Δmfe* | 0.22 ± 0.06 | 0.44 ± 0.09 | 16.76 ± 0.73 | 9.11 ± 0.33 | 0.35 ± 0.01 | 0.88 ± 0.05 | 11.36 ± 0.51 | 37.68 ± 0.69 | 20.03 ± 0.45 | 0.27 ± 0.02 | 0.69 ± 0.03 | 0.49 ± 0.03 | 0.61 ± 0.03 | - | 1.12 ± 0.04 | 31.20 ± 1.42 | 47.28 ± 1.03 | 21.52 ± 0.39 | 1.73 ± 0.06 ** |
| *Δmfe_acl* | 0.85 ± 0.10 | 0.24 ± 0.01 | 18.11 ± 0.47 | 11.41 ± 0.19 | 0.26 ± 0.03 | 0.79 ± 0.02 | 9.03 ± 0.25 | 37.38 ± 0.06 | 20.17 ± 0.58 | - | - | - | - | - | 1.77 ± 0.06 | 29.99 ± 0.62 | 48.79 ± 0.14 ** | 21.23 ± 0.54 | 1.77 ± 0.06 ** |
| *Δmfe_acc* | 0.30 ± 0.02 | 0.18 ± 0.01 | 14.81 ± 0.35 | 12.58 ± 0.39 | 0.25 ± 0.01 | 0.68 ± 0.03 | 6.46 ± 0.34 | 41.89 ± 0.28 | 21.67 ± 0.20 | - | - | - | - | - | 1.21 ± 0.04 | 22.94 ± 0.57 ** | 54.47 ± 0.45 ** | 22.59 ± 0.17 | 1.21 ± 0.04 ** |
| *Δmfe_ts* | 0.62 ± 0.03 | 0.24 ± 0.01 | 14.67 ± 0.54 | 12.93 ± 0.16 | 0.12 ± 0.01 | 0.69 ± 0.01 | 6.60 ± 0.24 | 47.83 ± 0.69 | 15 ± 0.43 | - | - | - | - | - | 1.30 ± 0.16 | 23.43 ± 0.30 ** | 60.76 ± 0.73 ** | 15.81 ± 0.43 ** | 1.30 ± 0.16 ** |
| *Δmfe_dga* | - | 0.27 ± 0.01 | 14.74 ± 0.14 | 6.64 ± 0.05 | 0.14 ± 0.00 | 0.53 ± 0.01 | 15.21 ± 0.10 | 50.16 ± 0.16 | 8.91 ± 0.11 | 0.14 ± 0.00 | 0.81 ± 0.03 | 0.53 ± 0.04 | 0.71 ± 0.01 | 0.27 ± 0.02 | 0.96 ± 0.06 | 32.71 ± 0.25 * | 57.60 ± 0.12 ** | 9.72 ± 0.11 ** | 1.94 ± 0.05 ** |
| *Δmfe_Δcex* | 0.44 ± 0.04 | 0.52 ± 0.04 | 16.95 ± 0.33 | 10.57 ± 0.15 | 0.28 ± 0.01 | 0.86 ± 0.05 | 10.86 ± 0.33 | 41.40 ± 0.68 | 14.85 ± 0.33 | - | 0.65 ± 0.03 | 0.59 ± 0.04 | 0.58 ± 0.01 | - | 1.46 ± 0.12 | 31.45 ± 0.75 | 52.56 ± 0.86 ** | 15.99 ± 0.32 ** | 2.04 ± 0.13 ** |
| *Δmfe_Δcex_acl* | 0.49 ± 0.13 | 0.32 ± 0.01 | 15.11 ± 0.71 | 11.79 ± 0.51 | 0.66 ± 0.05 | 0.73 ± 0.03 | 5.71 ± 0.26 | 41.97 ± 0.84 | 19.51 ± 0.36 | 0.58 ± 0.00 | - | 0.61 ± 0.02 | 0.31 ± 0.01 | - | 2.20 ± 0.13 | 24.15 ± 0.35 ** | 54.37 ± 0.51 ** | 21.48 ± 0.40 | 2.51 ± 0.13 |
| *Δmfe_Δcex_acc* | 0.41 ± 0.00 | 0.36 ± 0.02 | 15.30 ± 0.29 | 10.94 ± 0.30 | 0.28 ± 0.01 | 0.69 ± 0.05 | 6.24 ± 0.58 | 43.54 ± 0.30 | 18.94 ± 0.12 | 0.56 ± 0.03 | - | 0.66 ± 0.08 | 0.30 ± 0.00 | - | 1.93 ± 0.09 | 24.53 ± 0.49 ** | 55.13 ± 0.52 ** | 20.47 ± 0.17 ** | 2.23 ± 0.10 * |
| *Δmfe_Δcex_ts* | 0.42 ± 0.02 | 0.36 ± 0.02 | 15.49 ± 0.05 | 10.90 ± 0.23 | 0.61 ± 0.02 | 0.71 ± 0.03 | 6.92 ± 0.16 | 42.13 ± 0.74 | 19.18 ± 0.52 | 0.53 ± 0.09 | - | 0.56 ± 0.01 | 0.30 ± 0.00 | - | 2.14 ± 0.13 | 25.62 ± 0.33 ** | 53.59 ± 0.54 ** | 21.02 ± 0.44 * | 2.44 ± 0.13 |
| *Δmfe_Δcex_dga* | - | 0.20 ± 0.01 | 16.39 ± 0.58 | 6.65 ± 0.29 | - | 0.33 ± 0.03 | 14.14 ± 0.53 | 51.65 ± 0.30 | 5.40 ± 0.30 | - | 0.85 ± 0.02 | 1.15 ± 0.04 | 0.76 ± 0.01 | 0.52 ± 0.06 | 1.91 ± 0.13 | 34.26 ± 0.23 ** | 59.97 ± 0.57 ** | 5.89 ± 0.31 ** | 3.19 ± 0.08 * |
| *Δmfe_Δcex_dga_ts* | - | 0.22 ± 0.01 | 15.15 ± 0.33 | 6.48 ± 0.23 | 0.17 ± 0.00 | 0.43 ± 0.01 | 14.50 ± 0.23 | 49.80 ± 0.44 | 8.71 ± 0.17 | 0.11 ± 0.01 | 1.05 ± 0.03 | 0.57 ± 0.02 | 0.80 ± 0.03 | 0.32 ± 0.01 | 1.79 ± 0.12 | 33.50 ± 0.58 * | 57.16 ± 0.59 ** | 9.42 ± 0.19 ** | 2.90 ± 0.14 |
| Palm (range)<br>(Montoya et al., 2013) | - |  | 32.6–46.0 | 0.0–0.5 | - | - | 1.7–7.6 | 35.1–53.4 | 7.0–15.0 | 0.2–0.6 | 0.1–0.5 | 0.1–0.2 | - |  | - | 36.7–51.5 | 35.3–53.8 | 7.3–15.5 |  |

Differences with control (WT) were evaluated using a t-test \*: $p \leq 0.05$ , \*\*: $p \leq 0.01$ .

**Table S7.**
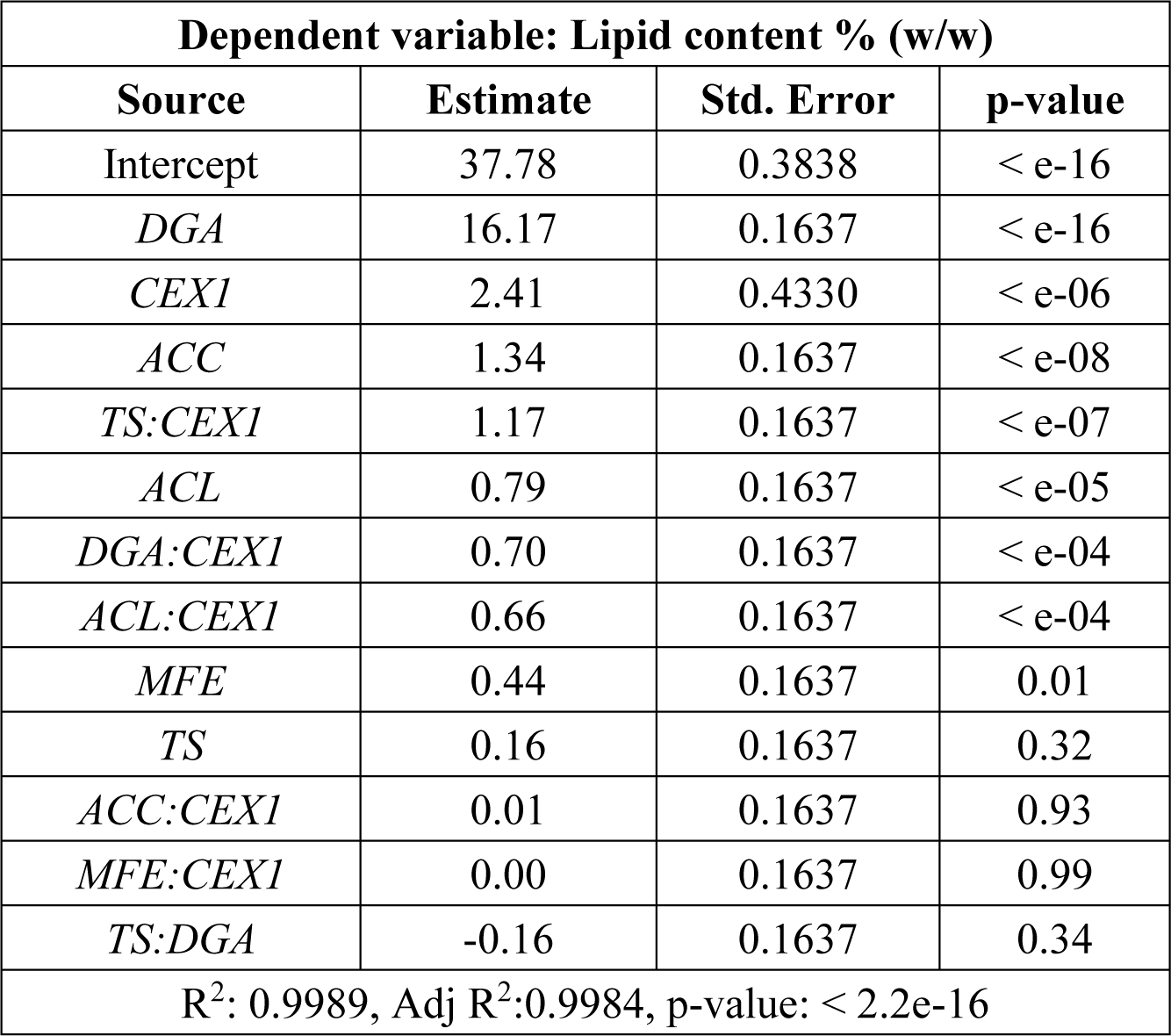
Model coefficients, and statistics of quadratic regression model.

## References

Abeln, F. and Chuck, C.J. (2021) The history, state of the art and future prospects for oleaginous yeast research. Microb Cell Fact 20: 221.

Abubakar, A., Ishak, M.Y., and Makmom, A.A. (2021) Impacts of and adaptation to climate change on the oil palm in Malaysia: a systematic review. Environmental Science and Pollution Research.

Amalia, L., Zhang, Y.H., Ju, Y.H., and Tsai, S.L. (2020) Enhanced Lipid Production in *Yarrowia lipolytica* Po1g by Over-expressing lro1 Gene under Two Different Promoters. Appl Biochem Biotechnol.

André, A., Chatzifragkou, A., Diamantopoulou, P., Sarris, D., Philippoussis, A., Galiotou-Panayotou, M., et al. (2009) Biotechnological conversions of bio-diesel-derived crude glycerol by *Yarrowia lipolytica* strains. Eng Life Sci 9: 468–478.

Beopoulos, A., Cescut, J., Haddouche, R., Uribelarrea, J.-L., Molina-Jouve, C., and Nicaud, J.-M. (2009) *Yarrowia lipolytica* as a model for bio-oil production. Prog Lipid Res 48: 375–387.

Beopoulos, A., Haddouche, R., Kabran, P., Dulermo, T., Chardot, T., and Nicaud, J.-M. (2012) Identification and characterization of DGA2, an acyltransferase of the DGAT1 acyl-CoA:diacylglycerol acyltransferase family in the oleaginous yeast *Yarrowia lipolytica*. New insights into the storage lipid metabolism of oleaginous yeasts. Appl Microbiol Biotechnol 93: 1523–1537.

Beopoulos, A., Nicaud, J.M., and Gaillardin, C. (2011) An overview of lipid metabolism in yeasts and its impact on biotechnological processes. Appl Microbiol Biotechnol.

Blazeck, J., Hill, A., Liu, L., Knight, R., Miller, J., Pan, A., et al. (2014) Harnessing *Yarrowia lipolytica* lipogenesis to create a platform for lipid and biofuel production. Nat Commun.

Blazeck, J., Liu, L., Knight, R., and Alper, H.S. (2013) Heterologous production of pentane in the oleaginous yeast *Yarrowia lipolytica*. J Biotechnol 165: 184–194.

Caporusso, A., Capece, A., and De Bari, I. (2021) Oleaginous Yeasts as Cell Factories for the Sustainable Production of Microbial Lipids by the Valorization of Agri-Food Wastes. Fermentation 7: 50.

Carsanba, E., Papanikolaou, S., and Erten, H. (2018) Production of oils and fats by oleaginous microorganisms with an emphasis given to the potential of the nonconventional yeast *Yarrowia lipolytica*. Crit Rev Biotechnol 38: 1230–1243.

Dobrowolski, A. and Mirończuk, A.M. (2020) The influence of transketolase on lipid biosynthesis in the yeast *Yarrowia lipolytica*. Microb Cell Fact 19: 138.

Dobrowolski, A., Mituła, P., Rymowicz, W., and Mirończuk, A.M. (2016) Efficient conversion of crude glycerol from various industrial wastes into single cell oil by yeast *Yarrowia lipolytica*. Bioresour Technol 207: 237–243.

Dulermo, T., Lazar, Z., Dulermo, R., Rakicka, M., Haddouche, R., and Nicaud, J.-M. (2015) Analysis of ATP-citrate lyase and malic enzyme mutants of *Yarrowia lipolytica* points out the importance of mannitol metabolism in fatty acid synthesis. Biochimica et Biophysica Acta (BBA) - Molecular and Cell Biology of Lipids 1851: 1107–1117.

Dulermo, T. and Nicaud, J.M. (2011) Involvement of the G3P shuttle and Β-oxidation pathway in the control of TAG synthesis and lipid accumulation in *Yarrowia lipolytica*. Metab Eng 13: 482–491.

Duman-Özdamar, Z.E., Julsing, M.K., Verbokkem, J.A.C., Wolbert, E., Martins dos Santos, V.A.P., Hugenholtz, J., and Suarez-Diez, M. (2024) Model-driven engineering of Cutaneotrichosporon oleaginosus ATCC 20509 for improved microbial oil production.

Duman-Özdamar, Z.E., Martins dos Santos, V.A.P., Hugenholtz, J., and Suarez-Diez, M. (2022) Tailoring and optimizing fatty acid production by oleaginous yeasts through the systematic exploration of their physiological fitness. Microb Cell Fact 21: 228.

Erian, A.M., Egermeier, M., Rassinger, A., Marx, H., and Sauer, M. (2020) Identification of the citrate exporter Cex1 of *Yarrowia lipolytica*. FEMS Yeast Res 20:.

Fabiszewska, A., Misiukiewicz-Stępień, P., Paplińska-Goryca, M., Zieniuk, B., and Białecka-Florjańczyk, E. (2019) An Insight into Storage Lipid Synthesis by *Yarrowia lipolytica* Yeast Relating to Lipid and Sugar Substrates Metabolism. Biomolecules 9: 685.

Fabiszewska, Misiukiewicz-Stępień, Paplińska-Goryca, Zieniuk, and Białecka-Florjańczyk (2019) An Insight into Storage Lipid Synthesis by *Yarrowia lipolytica* Yeast Relating to Lipid and Sugar Substrates Metabolism. Biomolecules 9: 685.

Friedlander, J., Tsakraklides, V., Kamineni, A., Greenhagen, E.H., Consiglio, A.L., MacEwen, K., et al. (2016) Engineering of a high lipid producing *Yarrowia lipolytica* strain. Biotechnol Biofuels.

Gajdoš, P., Nicaud, J., and Čertík, M. (2017) Glycerol conversion into a single cell oil by engineered *Yarrowia lipolytica*. Eng Life Sci 17: 325–332.

Holley, R.A. and Patel, D. (2005) Improvement in shelf-life and safety of perishable foods by plant essential oils and smoke antimicrobials. Food Microbiol.

Kerkhoven, E.J., Pomraning, K.R., Baker, S.E., and Nielsen, J. (2016) Regulation of amino-acid metabolism controls flux to lipid accumulation in *Yarrowia lipolytica*. NPJ Syst Biol Appl 2: 16005.

Kim, M., Park, B.G., Kim, E.-J., Kim, J., and Kim, B.-G. (2019) In silico identification of metabolic engineering strategies for improved lipid production in *Yarrowia lipolytica* by genome-scale metabolic modeling. Biotechnol Biofuels 12: 187.

Larroude, M., Park, Y., Soudier, P., Kubiak, M., Nicaud, J., and Rossignol, T. (2019) A modular Golden Gate toolkit for *Yarrowia lipolytica* synthetic biology. Microb Biotechnol 12: 1249–1259.

Lazar, Z., Neuvéglise, C., Rossignol, T., Devillers, H., Morin, N., Robak, M., et al. (2017) Characterization of hexose transporters in *Yarrowia lipolytica* reveals new groups of Sugar Porters involved in yeast growth. Fungal Genetics and Biology 100: 1–12.

Liu, H., Song, Y., Fan, X., Wang, C., Lu, X., and Tian, Y. (2021) *Yarrowia lipolytica* as an Oleaginous Platform for the Production of Value-Added Fatty Acid-Based Bioproducts. Front Microbiol 11:.

Moeller, L., Grünberg, M., Zehnsdorf, A., Aurich, A., Bley, T., and Strehlitz, B. (2011) Repeated fed-batch fermentation using biosensor online control for citric acid production by *Yarrowia lipolytica*. J Biotechnol 153: 133–137.

Montoya, C., Lopes, R., Flori, A., Cros, D., Cuellar, T., Summo, M., et al. (2013) Quantitative trait loci (QTLs) analysis of palm oil fatty acid composition in an interspecific pseudo-backcross from Elaeis oleifera (H.B.K.) Cortés and oil palm (Elaeis guineensis Jacq.). Tree Genet Genomes 9: 1207–1225.

Murphy, D.J., Goggin, K., and Paterson, R.R.M. (2021) Oil palm in the 2020s and beyond: challenges and solutions. CABI Agriculture and Bioscience 2: 39.

Odoni, D.I., Vazquez-Vilar, M., van Gaal, M.P., Schonewille, T., Martins dos Santos, V.A.P., Tamayo-Ramos, J.A., et al. (2019) *Aspergillus niger* citrate exporter revealed by comparison of two alternative citrate producing conditions. FEMS Microbiol Lett 366:.

P. Desbois, A. (2012) Potential Applications of Antimicrobial Fatty Acids in Medicine, Agriculture and Other Industries. Recent Pat Antiinfect Drug Discov.

Papanikolaou, S. and Aggelis, G. (2003) Modeling Lipid Accumulation and Degradation in *Yarrowia lipolytica* Cultivated on Industrial Fats. Curr Microbiol 46: 398–402.

Park, Y.K., Ledesma-Amaro, R., and Nicaud, J.M. (2020) De novo Biosynthesis of Odd-Chain Fatty Acids in *Yarrowia lipolytica* Enabled by Modular Pathway Engineering. Front Bioeng Biotechnol.

Poontawee, R., Lorliam, W., Polburee, P., and Limtong, S. (2023) Oleaginous yeasts: Biodiversity and cultivation. Fungal Biol Rev 44: 100295.

R Core Team (2020) R software: Version 4.0.2. R Foundation for Statistical Computing.

Ratledge, C. and Wynn, J.P. (2002) The biochemistry and molecular biology of lipid accumulation in oleaginous microorganisms. In Advances in Applied Microbiology.

van Rosmalen, R.P., Moreno-Paz, S., Duman-Özdamar, Z.E., and Suarez-Diez, M. (2024) CFSA: Comparative flux sampling analysis as a guide for strain design. Metab Eng Commun 19: e00244.

Rustan, A.C. and Drevon, C.A. (2005) Fatty Acids: Structures and Properties. In Encyclopedia of Life Sciences.

Rywińska, A., Juszczyk, P., Wojtatowicz, M., Robak, M., Lazar, Z., Tomaszewska, L., and Rymowicz, W. (2013) Glycerol as a promising substrate for *Yarrowia lipolytica* biotechnological applications. Biomass Bioenergy.

Sagnak, R., Cochot, S., Molina-Jouve, C., Nicaud, J.-M., and Guillouet, S.E. (2018) Modulation of the Glycerol Phosphate availability led to concomitant reduction in the citric acid excretion and increase in lipid content and yield in *Yarrowia lipolytica*. J Biotechnol 265: 40–45.

Salvador López, J.M., Vandeputte, M., and Van Bogaert, I.N.A. (2022) Oleaginous yeasts: Time to rethink the definition? Yeast 39: 553–606.

Sara, M., Brar, S.K., and Blais, J.F. (2016) Lipid production by *Yarrowia lipolytica* grown on biodiesel-derived crude glycerol: optimization of growth parameters and their effects on the fermentation efficiency. RSC Adv 6: 90547–90558.

Sitepu, I.R., Garay, L.A., Sestric, R., Levin, D., Block, D.E., German, J.B., and Boundy-Mills, K.L. (2014) Oleaginous yeasts for biodiesel: Current and future trends in biology and production. Biotechnol Adv 32: 1336–1360.

Tai, M. and Stephanopoulos, G. (2013) Engineering the push and pull of lipid biosynthesis in oleaginous yeast *Yarrowia lipolytica* for biofuel production. Metab Eng.

Tsirigka, A., Theodosiou, E., Patsios, S.I., Tsoureki, A., Andreadelli, A., Papa, E., et al. (2023) Novel evolved *Yarrowia lipolytica* strains for enhanced growth and lipid content under high concentrations of crude glycerol. Microb Cell Fact 22: 62.

Vijay, V., Pimm, S.L., Jenkins, C.N., and Smith, S.J. (2016) The Impacts of Oil Palm on Recent Deforestation and Biodiversity Loss. PLoS One 11: e0159668.

Wang, J., Ledesma-Amaro, R., Wei, Y., Ji, B., and Ji, X.-J. (2020) Metabolic engineering for increased lipid accumulation in *Yarrowia lipolytica* – A Review. Bioresour Technol 313: 123707.

Wasylenko, T.M., Ahn, W.S., and Stephanopoulos, G. (2015) The oxidative pentose phosphate pathway is the primary source of NADPH for lipid overproduction from glucose in *Yarrowia lipolytica*. Metab Eng 30: 27–39.

Wei, S., Jian, X., Chen, J., Zhang, C., and Hua, Q. (2017) Reconstruction of genome-scale metabolic model of Yarrowia lipolytica and its application in overproduction of triacylglycerol. Bioresour Bioprocess 4:.

Wen, Z. and Al Makishah, N.H. (2022) Recent advances in genetic technology development of oleaginous yeasts. Appl Microbiol Biotechnol.

Zhang, H., Zhang, L., Chen, H., Chen, Y.Q., Chen, W., Song, Y., and Ratledge, C. (2014) Enhanced lipid accumulation in the yeast *Yarrowia lipolytica* by over-expression of ATP: Citrate lyase from Mus musculus. J Biotechnol 192: 78–84.

Zinjarde, S.S. (2014) Food-related applications of *Yarrowia lipolytica*. Food Chem.

